# Transcriptional Circuitry in HGSOC: A Dynamic Three-State Model Informed by a Living Biobank of Purified Tumour Fractions

**DOI:** 10.1101/2025.07.18.665513

**Authors:** Bethany M. Barnes, Samantha Littler, Anthony Tighe, Alicia Evans, James Altringham, Louisa Nelson, I-Hsuan Lin, Robert D. Morgan, Joanne C. McGrail, Stephen S. Taylor

## Abstract

High-grade serous ovarian cancer (HGSOC) is a heterogeneous disease, but efforts to define transcriptional subtypes using bulk RNA sequencing have been confounded by the presence of non-malignant cells. As a result, it remains unclear whether tumour-cell-intrinsic states exist, and whether these represent stable disease subtypes or are dynamically remodelled during disease progression and treatment. Here, we address this question using a living biobank of patient-derived ovarian cancer models (OCMs) cultured as purified tumour-cell populations under uniform conditions. RNA sequencing followed by unsupervised non-negative matrix factorisation (NMF) revealed a robust, hierarchical architecture comprising three core tumour-cell-intrinsic subtypes: the ***Alpha*** cluster, marked by cell-cycle deregulation and E2F-driven replication stress; the ***Beta*** cluster, defined by tumour-cell-intrinsic immune mimicry and inflammatory signalling; and the ***Gamma*** cluster, characterised by epithelial identity, extracellular matrix engagement, and metabolic adaptation. At higher clustering resolution, a fourth cluster, ***Delta***, emerged as a ***Gamma*** sub-lineage distinguished by a vesicle-oriented, neuronal-like secretory programme. By projecting cluster labels onto a subset of matched longitudinal OCMs using non-negative least squares, we show that while some tumours retain stable subtype identities, others display transcriptional plasticity, including transitions from epithelial-like ***Gamma*** states to more proliferative or secretory phenotypes. Together, these findings define the core architecture and dynamic potential of tumour-cell-intrinsic transcriptional states within HGSOC, thereby bridging legacy bulk classifications with emerging single-cell insights, establishing a framework for more precise patient stratification.

## INTRODUCTION

Epithelial ovarian cancer is an umbrella term for five distinct histological diseases—high-grade serous, low-grade serous, endometrioid, clear cell, and mucinous—each with distinct cells of origin, molecular alterations, and treatment responses (WHO Classification of Tumours Editorial Board, 2020). High-grade serous ovarian carcinoma (HGSOC) is characterised by near-universal *TP53* mutations and profound chromosomal instability (TCGA, 2011). By contrast, low-grade serous ovarian carcinomas (LGSOC) maintain relative genomic stability but commonly show RAS–RAF–MAPK pathway mutations (Singer et al., 2005; Etemadmoghadam et al., 2017). Endometrioid and clear cell ovarian cancers are often associated with endometriosis and frequently carry *ARID1A* or *PIK3CA* mutations (Wiegand et al., 2010; Friedlander et al., 2016). Mucinous ovarian cancer is distinct again, with *KRAS* mutations and molecular features reminiscent of gastrointestinal neoplasms (Cheasley et al., 2019). These differences are reflected in characteristic gene expression patterns that can distinguish histological subtypes and deepen our understanding of their unique biology (Domcke et al., 2013; Barnes et al., 2021).

Among these subtypes, HGSOC accounts for approximately 70–80% of epithelial ovarian cancer cases and is characterised by rapid clinical progression and frequent late-stage diagnosis (Gaitskell et al., 2022; Narayanan et al., 2025). The introduction of homologous recombination deficiency (HRD) biomarkers has improved treatment precision by identifying patients likely to benefit from PARP inhibitors (Coleman et al., 2019; Gonzalez-Martin et al., 2019; Ray-Coquard et al., 2019; Monk et al., 2022). However, because only 40– 50% of HGSOC cases are HRD-positive, this approach leaves a substantial fraction of patients without a clear biomarker-guided option (Morgan et al., 2023). This gap underscores a fundamental question: does HGSOC harbour transcriptional subtypes that could expose new therapeutic vulnerabilities?

Early answers to this question came from landmark studies that analysed gene expression in bulk tumours to classify HGSOC. Tothill et al. (2008) first defined four gene expression subtypes in HGSOC, a structure that TCGA (2011) and other studies have broadly confirmed (Helland et al., 2011; Verhaak et al., 2013; Konecny et al., 2014; Tan et al., 2015; Way et al., 2016; Chen et al., 2018; Talhouk et al., 2020). The C1/Mesenchymal subtype is defined by a desmoplastic stroma, high extracellular matrix (ECM) gene expression, and poorer prognosis. The C2/Immunoreactive group shows strong T-cell infiltration and inflammatory signalling, generally linked to improved survival. The C4/Differentiated subtype retains high epithelial marker expression (e.g., *MUC16*) and moderate immune signals. By contrast, the C5/Proliferative cluster is depleted of stromal and immune elements and is driven by upregulation of cell-cycle regulators and stem-like programs, correlating with aggressive clinical behaviour. Together, these studies showed that gene expression captures biologically and clinically relevant features in HGSOC that carry prognostic weight.

Emerging evidence from single-cell RNA sequencing (scRNA-seq), spatial transcriptomics, and laser-capture microdissection has demonstrated that the C1/Mesenchymal and C2/Immunoreactive transcriptional subtypes of HGSOC predominantly arise from non-malignant components of the tumour microenvironment.

Specifically, scRNA-seq studies (Izar et al., 2020; Olbrecht et al., 2021; Deng et al., 2022) consistently demonstrate that the C1/Mesenchymal subtype localises to cancer-associated fibroblasts and myofibroblasts, while the C2/Immunoreactive subtype is driven by T cells, NK cells, dendritic cells, and myeloid populations, rather than transcriptional programmes intrinsic to the tumour cells themselves. Spatial profiling (Denisenko et al., 2024) and laser-capture microdissection (Schwede et al., 2020) confirm that these signals are enriched in stromal and immune-rich regions but diminish when analysing pure tumour-cell areas. In contrast, the C4/Differentiated and C5/Proliferative subtypes appear to represent *bona fide* tumour-cell-intrinsic subtypes: scRNA-seq indicates that malignant epithelial cells segregate into roughly mutually exclusive populations with either high C5/Proliferative activity or high C4/Differentiated scores (Izar et al., 2020). This mutual exclusivity supports the interpretation that these subtypes reflect tumour-cell-intrinsic transcriptional subtypes that likely remain detectable in bulk RNA profiling due to minimal confounding by stromal or immune cell admixture.

These converging findings expose a persistent blindspot: bulk profiling reproducibly uncovers transcriptional clusters, but multiple lines of evidence, together with single-cell and spatial data, show that two of the four classical subtypes are driven by extrinsic cell types. We do not know whether the remaining two subtypes fully reflect the spectrum of heterogeneity in HGSOC, or whether additional subtypes are obscured by bulk tumour analysis. And if true tumour-cell-intrinsic subtypes exist in HGSOC, are these stable disease states or dynamic phenotypes shaped by the microenvironment, therapy, and tumour evolution? Answering this requires resolution and scale, a combination that current single-cell and spatial methods, while powerful, cannot yet deliver affordably at scale.

To revisit these questions from a new angle, we leveraged a living biobank of patient-derived ovarian cancer models (OCMs), *ex vivo* cultures of purified tumour cells grown in defined medium (Nelson et al., 2020; Barnes et al., 2021; Nelson et al., 2023). We performed RNA sequencing (RNA-seq) on 88 OCMs from 70 patients with histologically confirmed HGSOC, including a subset of longitudinal models collected across different stages of treatment. We applied an unsupervised machine-learning method (non-negative matrix factorisation [NMF]) to resolve transcriptional subtypes. To explore whether these clusters represent stable subtypes or dynamic states, we then projected longitudinal OCMs onto the cluster framework using non-negative least squares (NNLS), quantifying how subtype identity shifts over time and treatment.

## RESULTS

### Molecular validation of patient-derived ovarian cancer models

To overcome the limitations of traditional ovarian cancer cell lines—which often lack clinical annotation and do not capture the full molecular heterogeneity of HGSOC—we established a living biobank of patient-derived OCMs (Nelson et al., 2020; Nelson et al., 2023). These models are developed from either ascitic fluid or solid tumour biopsies collected during routine care, and are clinically annotated with detailed treatment histories and patient outcome data (**Figure 1a**). Importantly, OCMs are amenable to molecular profiling, including RNA-seq, functional interrogation and drug screening (Pillay et al., 2019; Coulson-Gilmer et al., 2021; Golder et al., 2022; Coulson-Gilmer et al., 2024; Littler et al., 2025; Tighe et al., 2025).

**Figure 1.**
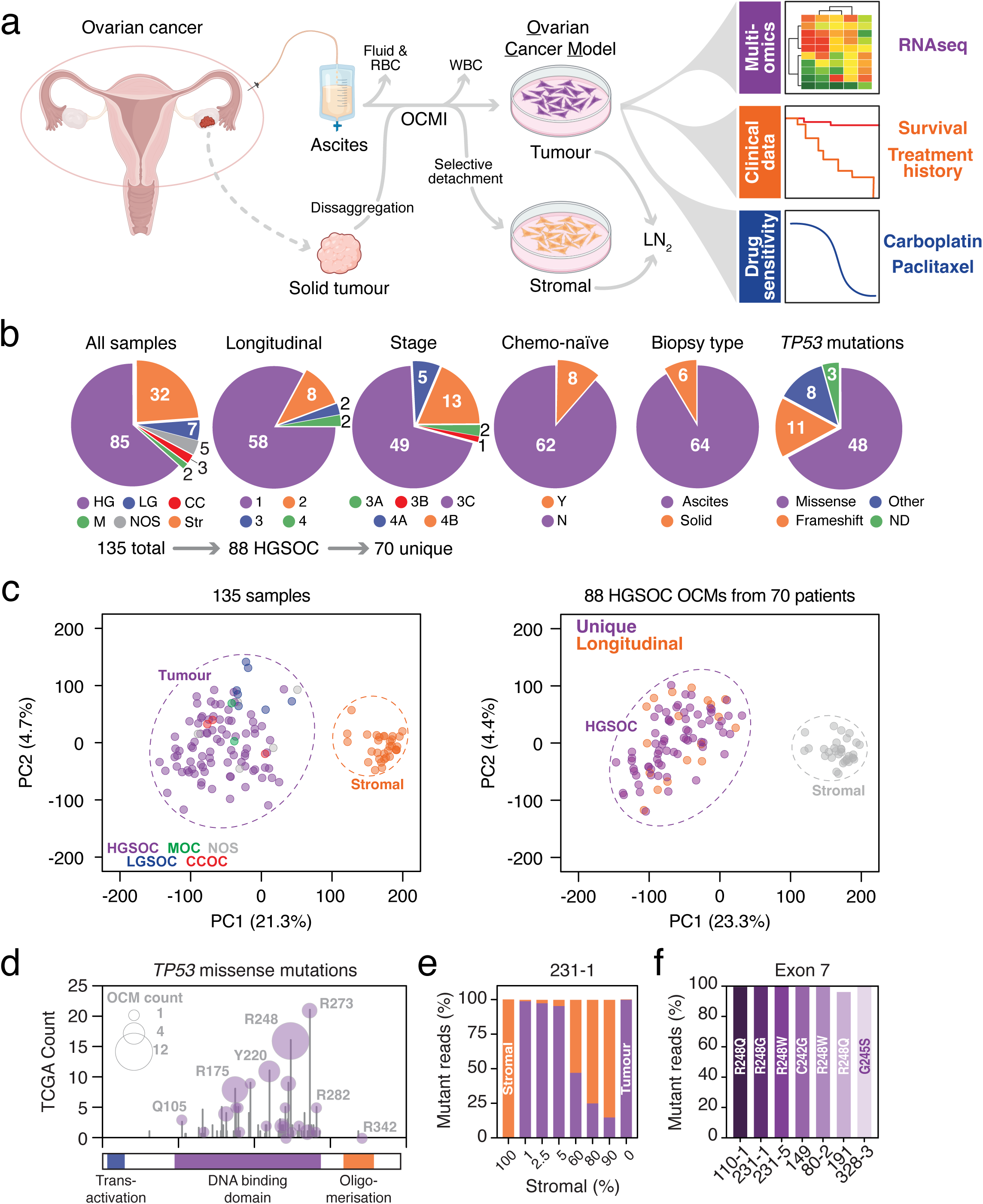
Molecular Validation of Patient-Derived Ovarian Cancer Models. (a) Overview of the OCM pipeline. Tumours were enzymatically dissociated; ascites were centrifuged and RBCs lysed. Cells were cultured in OCMI medium. Stromal fibroblasts were depleted through serial trypsinisation, yielding purified tumour cell cultures. OCMs were cryopreserved and expanded for RNA sequencing and drug screening. Created in BioRender. Taylor, S. (2025) https://BioRender.com/ohdtdas (b) Clinical and molecular features of the cohort. From 103 tumour-purified cultures and 32 stromal cultures, 88 HGSOC OCMs from 70 patients were retained after excluding non-HGSOC or reclassified samples (Barnes et al., 2021). Pie charts show cohort attributes: longitudinal sampling, FIGO stage, prior chemotherapy, biopsy site, and *TP53* mutation status. (c) PCA of transcriptomes. Left: All OCMs and stromal fibroblast cultures coloured by histopathology. Dashed ellipses highlight tumour vs. stromal separation. Right: Final HGSOC cohort, coloured by longitudinal vs. unique sampling; stromal fibroblast cultures shown in grey. (d) Analysis of the *TP53* missense mutations in the living biobank. Circle size reflects OCM count per mutation; stem height indicates frequency in TCGA dataset of HGSOC samples. Mutations are mapped along *TP53* domains. (e) *TP53* exon 7 sequencing of OCM.231-1 mixed at defined tumour:stromal ratios shows declining mutant allele frequency with increasing stromal content. (f) *TP53* Exon 7 sequencing of seven representative OCMs. Abbreviations: RBC, red blood cell; LN₂, liquid nitrogen; HG, high-grade serous; LG, low-grade serous; CC, clear cell; M, mucinous; NOS, not otherwise specified; Str, stromal; WT, wild-type; ND, not determined.

To date, RNA-seq has been performed on 135 patient-derived samples (**Figure 1b, c**), including 103 OCMs and 32 matched stromal fibroblast cultures. Based on histopathology, we excluded 12 OCMs derived from non-HGSOC subtypes. An additional five OCMs were derived from ovarian adenocarcinomas with unspecified subtypes. Using *TP53* mutation status and a previously described transcriptomic classifier (Barnes et al., 2021), two of these were excluded as likely LGSOC-derived, while the remaining three, which display hallmarks of HGSOC, were retained. This resulted in a final cohort of 88 HGSOC-derived OCMs (**Figure 1b, c**). These models were derived from 70 patients, 12 of whom contributed longitudinal biopsies that gave rise to additional models. At diagnosis, all patients presented with FIGO stage IIIA–IVB disease (**Figure 1b** and **Table S1**). Most OCMs were established from ascitic fluid, with the remainder from solid tumour biopsies, and while eight models were chemo-naïve, most were derived from post-treatment samples. Sequencing cloned transcripts confirmed that all the selected HGSOC-derived OCMs harboured *TP53* mutations (**Figure 1b** and **Table S1**), consistent with the near-universal p53 disruption in this disease (Ahmed et al., 2010). *TP53* mutations included 48 missense mutations, predominantly in exons 5–8 of *TP53*, which encode the DNA-binding domain (**Figure 1d**), consistent with mutation hotspots observed in the TCGA HGSOC cohort (TCGA, 2011).

OCMs are highly purified tumour cell fractions, and accordingly, principal component analysis (PCA) of the RNA-seq data revealed clear separation between the OCMs and stromal fibroblasts (**Figure 1c**). To further test the purity of the OCMs, we performed amplicon-sequencing, targeting exons 5–7 of the *TP53* locus. Given that *TP53* mutation is a truncal event in HGSOC (Ahmed et al., 2010), the mutant allele frequency (MAF) should approach 100% in cultures composed exclusively of malignant cells. To validate the sensitivity of the sequencing assay, we spiked OCMs with increasing amounts of patient-matched stromal fibroblasts (**Figure 1e**). The MAF closely tracked with tumour cell content, and importantly, wild-type *TP53* alleles were detectable even when stromal cells comprised just 1% of the mixture. We then applied the assay to the OCMs, which revealed *TP53* MAFs of ≥96%, consistent with near-complete tumour cell purity and retention of the clonal *TP53* mutation. Together, these findings indicate that the OCMs are indeed highly purified, HGSOC-derived tumour cell fractions, establishing them as a robust platform for interrogating tumour-cell-intrinsic transcriptional states in HGSOC.

### Subtypes emerge from NMF clustering of HGSOC transcriptomes

To investigate the presence of transcriptional subtypes within HGSOC, we applied NMF to RNA-seq profiles from the earliest available OCM for each of the 70 patients. Recognising that clustering results can be sensitive to input gene selection, we systematically varied the number of genes used—generating ten input matrices comprising the top 1,000 to 10,000 most variable genes (based on median absolute deviation)—and assessed clustering performance across a range of ranks (, = 2 to 10). Clustering quality metrics, including cophenetic correlation coefficient (Brunet et al., 2004), dispersion (Kim and Park, 2007), and silhouette score (Rousseeuw, 1987), were remarkably consistent across gene list sizes, with peak values observed at *k* = 2 and *k* = 3, indicating that these ranks provided the most stable and well-separated solutions (**Figure 2a**). To examine these results in greater depth, we repeated NMF at *k* = 2 and *k* = 3 with 200 iterations across all ten gene input sets. Among these, the 3,000-gene input at *k* = 2 and the 6,000-gene input at *k* = 3 yielded the highest cophenetic coefficients (**Figure S1a**). These were selected for consensus matrix visualisation, showing how frequently each pair of samples co-clustered across NMF iterations (**Figures 2b** and **S1b**). The consensus matrix at *k*, = 3 (**Figure 2b**) revealed three well-defined clusters with strong pairwise co-clustering and clear separation, and most samples exhibited high silhouette widths, supporting confident cluster assignment.

**Figure 2.**
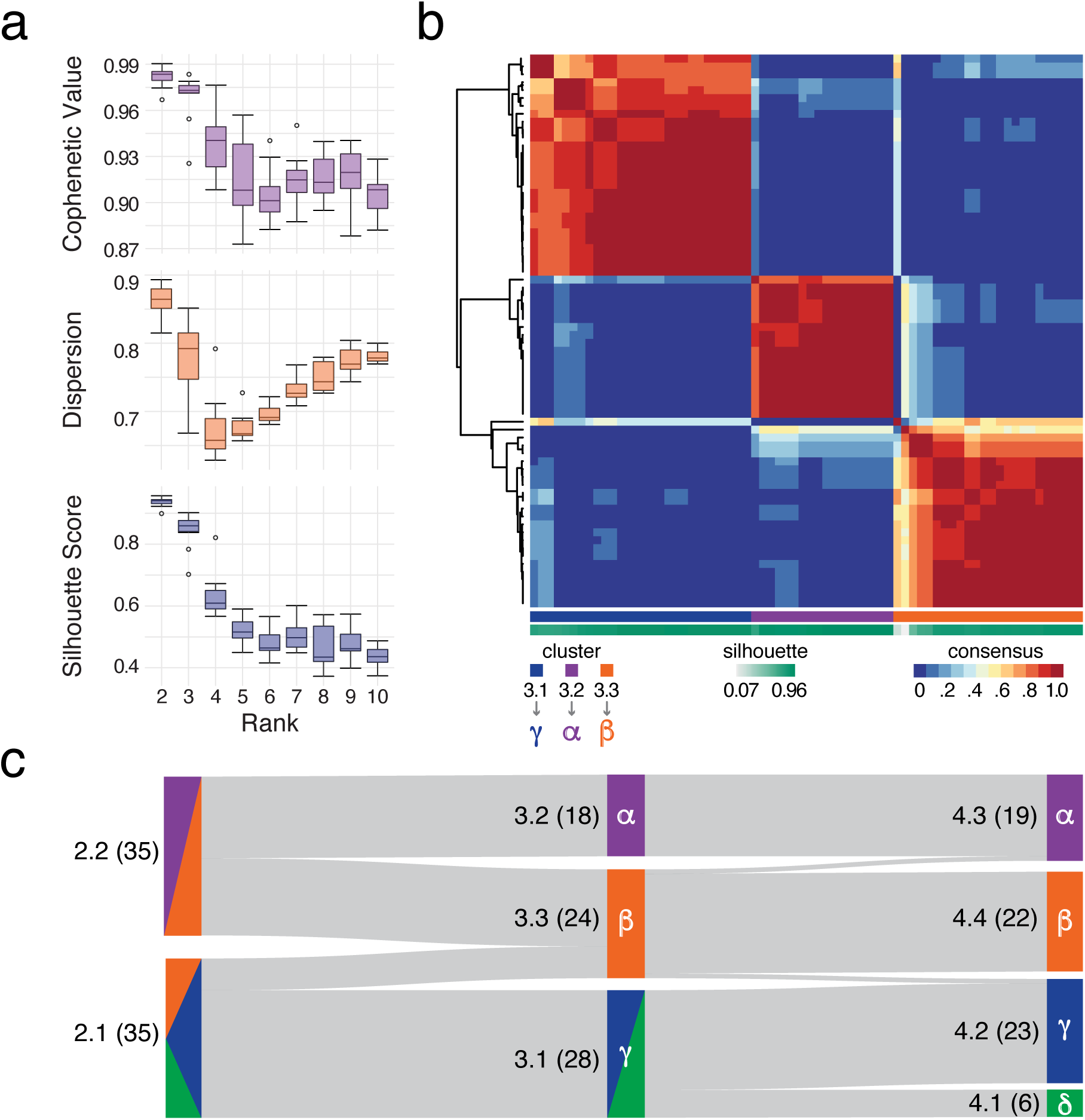
Subtypes emerge from NMF clustering of HGSOC transcriptomes. (a) Evaluation of NMF clustering performance across gene list sizes and cluster ranks. NMF clustering was applied to RNA-seq data from 70 ovarian cancer models (OCMs) using gene sets comprising the top 1,000 to 10,000 most variable genes. For each gene set, clustering was performed across ranks *k* = 2 to *k* = 10 (50 runs per condition). Clustering quality was evaluated using three metrics: cophenetic correlation (top), dispersion coefficient (middle), and silhouette score (bottom). Each boxplot summarises metric values across the ten gene sets. (b) Consensus clustering heatmap of 70 OCMs using the top 6,000 most variable genes and *k* = 3 clusters. The consensus matrix shows the pairwise frequency with which models co-cluster across 200 NMF iterations, with red indicating samples that consistently clustered together and blue indicating infrequent co-clustering. Dendrograms reflect hierarchical clustering of the consensus matrix. Bottom bars indicate NMF cluster assignment (top) and silhouette width for each sample (bottom). (c) Sankey plot illustrating the stability and evolution of sample cluster assignments across NMF ranks 2 to 4. For each sample, the dominant cluster assignment across gene input sizes was used at each rank. Numbers in parentheses indicate the number of OCMs in that subcluster.

To identify the clustering solution that best captured a biologically meaningful substructure, we examined how OCM assignments transitioned across cluster ranks using a Sankey plot (**Figure 2c**), based on each OCM’s modal cluster membership across gene list inputs (**Figure S1c**). At *k* = 2, samples segregated into two major groups, 2.1 and 2.2. Increasing to *k* = 3 revealed that most OCMs from cluster 2.1 transitioned to cluster 3.1, while most from 2.2 transitioned to cluster 3.2. Notably, a subset of 2.1 and 2.2 samples also transitioned into 3.3, such that 3.3 represents a recombination of samples from both *k* = 2 clusters. This partial mixing suggests that 3.3 captures a transcriptionally distinct subgroup not resolved at *k* = 2. The consensus matrix at *k* = 3 also revealed tightly co-clustered substructures within each cluster, particularly within 3.1 and 3.3, hinting at the presence of additional internal heterogeneity. To investigate this, we performed NMF at *k* = 4 (200 iterations across all gene lists) and mapped the resulting cluster assignments (**Figure S1c**) onto the Sankey plot (**Figure 2c**). At this resolution, clusters 3.2 and 3.3 remained highly stable, while 3.1 consistently divided into two subgroups, 4.1 and 4.2. This supports a hierarchical model in which 3.1 contains finer-grained transcriptional heterogeneity.

Heatmap visualisation of the dominant cluster assignment for each sample across input gene list sizes (**Figure S1c**) confirmed that cluster 4.1 reproducibly emerged as a stable subcluster of 3.1 across most gene sets, particularly at lower and mid-range input sizes. However, at larger input sizes (8,000–10,000 genes), an alternative pattern emerged in which cluster 3.3, rather than 3.1, appeared to subdivide—suggesting that additional finer-grained structures may be present. Despite this, the subdivision of 3.1 into two consistent subclusters was more frequently observed and better supported across gene list sizes, indicating a more stable and reproducible solution. These findings reinforce the robustness of the three core subtype structure, while also highlighting that further heterogeneity may be uncovered at higher clustering resolution. To facilitate further description, we assigned Greek symbols to the clusters, with *Alpha* (⍺), *Beta* (/), *Gamma* (0), and *Delta* (1) corresponding to clusters 3.2/4.3, 3.3/4.4, 3.1/4.2, and 4.1, respectively (**Figure 2c**).

### Characterisation of three core transcriptional subtypes

Having established three core transcriptional subtypes, we next investigated their defining molecular features. Differential gene expression analysis was performed using *limma-voom*, with each OCM assigned to its dominant cluster at *k* = 3. To account for variability in cluster assignments across input gene sets, we derived sample-specific weights based on cluster consistency (**Figure S1c**) and incorporated these into the linear model. For each subtype, gene expression was compared against the other two clusters combined, identifying 1,638 upregulated and 1,780 downregulated genes in *Alpha*; 601 upregulated and 848 downregulated in *Beta*; and 2,208 upregulated and 1,715 downregulated in *Gamma* (FDR < 0.05; **Figure S2a**). Pathway enrichment of upregulated genes was performed using g:Profiler2 across multiple databases, with manual curation of KEGG, Reactome, and selected Gene Ontology (GO) terms into consolidated functional themes (**Figure 3a** and **Table S2**). This analysis revealed a clear pattern: *Alpha* was dominated by cell cycle–related processes, *Beta* by inflammatory-related features, and *Gamma* by epithelial-associated biology. Below, we examine the molecular features of each core transcriptional subtype in greater detail.

**Figure 3.**
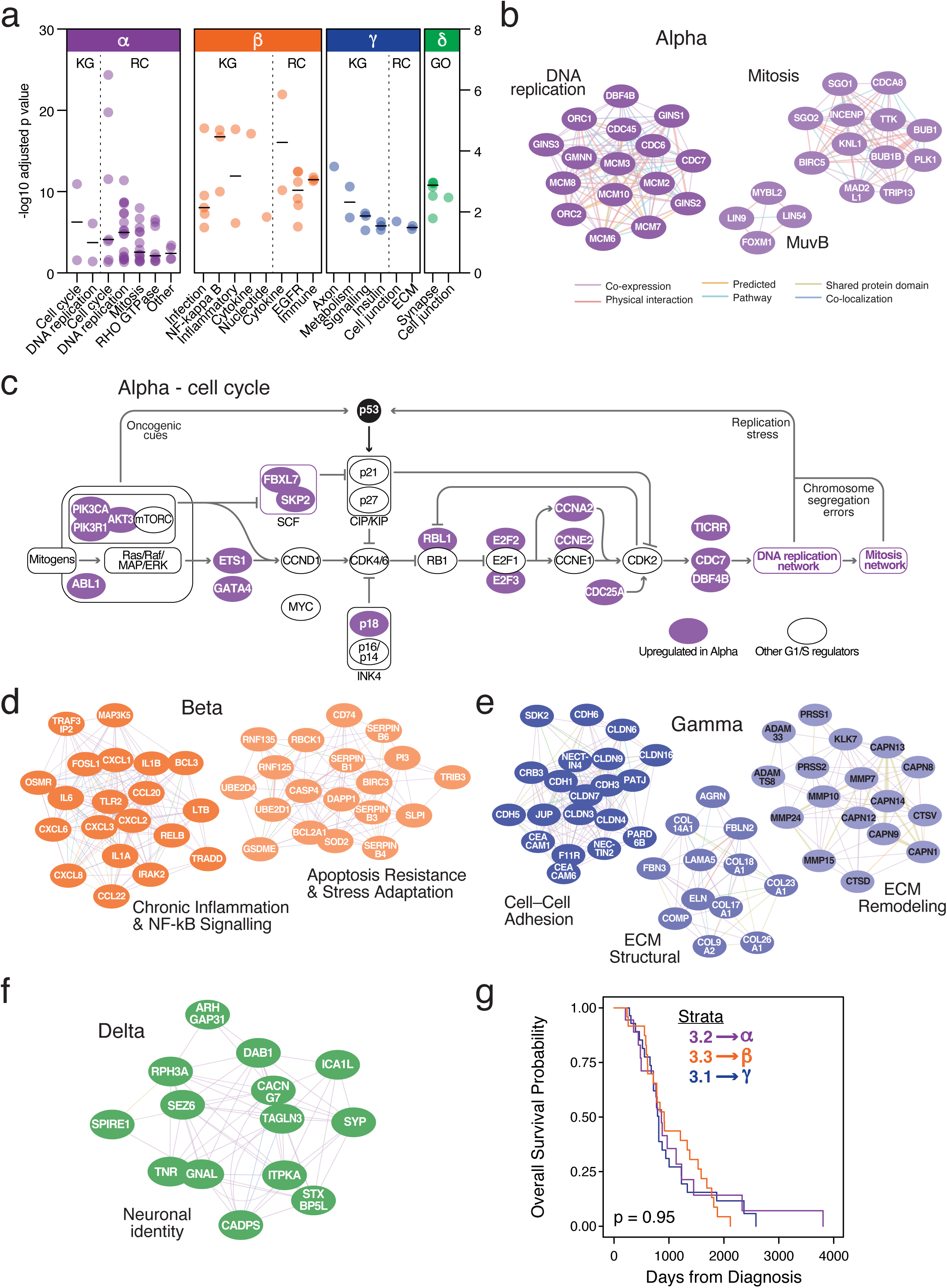
Characterisation of Transcriptional Subtypes. (a) KEGG (KG) and Reactome (RC) pathway enrichment of upregulated genes in transcriptional subtypes *Alpha* (purple), *Beta* (orange), and *Gamma* (blue) from the 3-cluster NMF solution, performed using g:Profiler2. Pathway terms were manually grouped into broader functional categories for clarity. Only statistically significant enrichments (FDR < 0.05) are shown. Gene Ontology (GO) enrichment for cluster *Delta* (green) is shown from the 4-cluster fit, where *Delta* was compared specifically with *Gamma*. (b) Network representation of key genes upregulated in subtype *Alpha*, generated using Cytoscape. Left: genes involved in DNA replication. Middle: members of the MuvB complex. Right: mitotic regulators. Edges indicate evidence of co-expression, physical interaction, shared domains, pathway co-membership, or co-localisation based on integrated annotation sources. (c) Schematic of the G1/S cell cycle transition pathway highlighting genes upregulated in subtype *Alpha* (purple). Upstream mitogenic and oncogenic signalling converges on RB/E2F axis deregulation, promoting CDK activation and progression through G1/S. Coordinated upregulation of E2F targets, DNA replication and mitotic genes reflects the transcriptional amplification of this programme. Grey nodes indicate canonical regulators not upregulated in *Alpha* . (d) Example networks of genes upregulated in subtype *Beta* (orange), highlighting modules associated with Chronic Inflammation & NF-κB Signalling and Apoptosis Resistance & Stress Adaptation. (e) Example network of genes upregulated in subtype *Gamma* (blue), grouped into three modules: Cell–Cell Adhesion, ECM Structural, and ECM Remodelling. (f) Example network of genes upregulated in subtype *Delta* (green): Neuronal Identity. (g) Kaplan–Meier survival analysis of transcriptional subtypes *Alpha*, *Beta*, and *Gamma* based on the 3-cluster NMF solution. Overall survival from diagnosis is plotted for each subtype, with no statistically significant difference observed (log-rank p = 0.095).

### Transcriptional subtype Alpha—cell cycle deregulation and proliferative signalling

Pathway enrichment analysis (KEGG and Reactome) for transcriptional subtype *Alpha* revealed multiple significantly enriched pathways, which were manually grouped into four overarching themes: G1/S control, DNA replication, mitotic signalling, and RHO-GTPase–mediated cytoskeletal regulation (**Figure 3a** and **Table S2**). Virtually every phase of the cell cycle was represented in subtype *Alpha* (**Figure 3b, c**). G1/S transition genes were upregulated, including *E2F2, E2F3, CCNE2, CCNA2, CDC25A, SKP2, CDKN2C,* and *RBL1*. Components of the DNA replication machinery—*CDC6, ORC1, ORC2, MCM2–7, CDC45, GINS1–3, POLA1, POLD1, POLE2, DNA2, FEN1,* and *EXO1*—were strongly expressed. G2/M progression and mitosis were marked by elevated expression of *PLK1, BUB1, BUB1B, TTK, MAD2L1, ANAPC1, CDCA8, INCENP, BIRC5, CENPF,* and *CENPU*, alongside activated MuvB complex components *MYBL2, LIN54, LIN9,* and *FOXM1*. Genes involved in the DNA damage response—including *CHEK1, CLSPN, BRCA2, BRIP1, MDC1,* and *BARD1*—as well as centrosome and spindle assembly genes such as *PLK4, PCNT, CEP152, CEP135, TPX2, KIF18A,* and *TUBA1B*– were also upregulated. In addition, enrichment of PI3K/AKT/mTOR, MAPK, and NOTCH signalling pathways suggests convergence of upstream mitogenic cues that may override cell cycle checkpoints (**Figure 3c** and **Table S2**).

TRANSFAC analysis further highlighted deregulated E2F signalling as a central feature, showing an enrichment of *E2F* target genes (**Table S2**). To confirm transcription factor (TF) activity, we applied CollecTRI—a curated meta-resource of signed TF–target interactions—which verified elevated activity of *E2F1-4* in subtype *Alpha* (**Figure S2c**). Subtype *Alpha* also expressed epithelial–mesenchymal transition (EMT) markers, including *TWIST1* and *TWIST2* (**Figure S2a**), along with components of the TGF-β and WNT signalling pathways (**Table S2**), suggesting potential acquisition of mesenchymal traits alongside proliferative deregulation.

In summary, transcriptional subtype *Alpha* is defined by a distinct upregulation of cell-cycle– associated genes, including those regulating DNA replication, mitosis, and various checkpoint controls. Alongside enrichment of DNA replication and repair pathways, this suggests a transcriptional programme primed for sustained proliferation under replicative stress

### Transcriptional subtype Beta—inflammatory signalling, immune modulation, and survival pathways

Pathway enrichment (KEGG and Reactome) for subtype *Beta* identified multiple upregulated pathways, which we manually grouped into infection response, cytokine and inflammatory signalling, immune regulation, nucleotide metabolism, and EGFR signalling (**Figure 3a** and **Table S2**). Elevated expression of a broad repertoire of pro-inflammatory cytokines and chemokines was evident, including *IL1A, IL1B, IL6, CXCL1, CXCL2, CXCL8,* and *CCL20*, as well as receptors and adaptors such as *TLR2, IL15RA, OSMR,* and *IL27RA* (**Figure 3d**). This profile suggests a heightened inflammatory state and supports the concept of immune mimicry, in which tumour cells adopt features of innate immune signalling. Additional immune-modulatory features included upregulation of antigen presentation genes (*CD74, ICAM3, CD40*) and immune checkpoint ligands such as *PDCD1LG2* (PD-L2), indicating potential strategies for immune engagement and evasion in an immunologically active microenvironment.

A central hallmark of subtype *Beta* was activation of the NF-κB pathway, supported by pathway enrichment and upregulation of upstream activators, including *TLR2, OSMR, CD40, IL15RA, IL27RA, IFNGR1, TNFRSF11B,* and *CD74*, which converge on canonical intermediates (*MyD88, TRADD, TRAFs*) to activate NF-κB. Correspondingly, downstream components (*TRADD, TRAF3IP2, RELB, FOSL1*) and NF-κB–responsive survival genes (*BIRC3, BCL2A1*) were also upregulated, indicating a tumour-cell-intrinsic inflammatory circuit. A related stress-adaptive survival module was also evident, highlighted by genes associated with apoptotic regulation and cellular stress responses (**Figure 3d**), including NF-κB targets (*BCL2A1, BIRC3, TRIB3, SLPI, SOD2*), pyroptosis-associated genes (*CASP4, GSDME*), protease inhibitors (*SERPINB1/B3/B4/B6*), and ubiquitin pathway regulators (*RNF125, RBCK1, UBE2D1/4*). Although not confined to a single curated pathway, this network supports the idea that *Beta* tumours are transcriptionally primed to tolerate stress and evade cell death.

In parallel, subtype *Beta* showed upregulation of multiple EGFR ligands (*AREG, HBEGF, TGFA, EPGN*) and EGFR itself. As EGFR signalling activates PI3K–AKT and MAPK pathways—both of which feed into NF-κB via IKK-mediated mechanisms—this suggests potential crosstalk and a positive feedback loop that reinforces inflammatory and pro-survival signalling. Several EGFR ligands are also known NF-κB transcriptional targets, further supporting this idea. Subtype *Beta* also showed significant enrichment of nucleotide metabolism pathways (**Figure 3a**), possible to support elevated biosynthetic demand and redox buffering in the context of inflammation and proliferation.

In summary, subtype *Beta* represents a transcriptionally distinct, tumour-cell-intrinsic inflammatory state marked by NF-κB activation, immune mimicry and modulation, and survival signalling. The convergence of mitogenic and inflammatory cues—via EGFR–NF-κB crosstalk—and the upregulation of stress tolerance programmes suggest a phenotype adapted for survival under immunological and therapeutic pressure.

### Transcriptional subtype Gamma—epithelial identity and ECM remodelling

Reactome enrichment analysis for subcluster *Gamma* prominently highlighted pathways related to cell–cell adhesion, tight junction assembly, and extracellular matrix (ECM) organisation (**Figure 3a** and **Table S2**). This was supported by coordinated upregulation of classical epithelial markers, including cadherins (*CDH1, CDH3*), claudins (*CLDN3, CLDN4, CLDN6*), and adhesion scaffolds (*NECTIN4, CRB3, PARD6B, PATJ*) (**Figure 3e**). Subtype *Gamma* also expressed canonical HGSOC markers (*PAX8, WT1, EPCAM, MUC16*), while mesenchymal and cancer stem cell–associated genes such as *CD44* and *VIM* were downregulated. Together, these features indicate a transcriptional state that retains epithelial identity and cell polarity, and one that is typically lost during EMT.

ECM remodelling and stromal engagement were also prominent, with high level expression of ECM components (*COL14A1, LAMA5, FBN3, ELN*) and remodelling enzymes, including matrix metalloproteinases (MMPs), cathepsins, and ADAM/ADAMTS family proteases. Integrins (*ITGA9, ITGB4/6/8*) and matrix receptors (*DDR1*) were also enriched, suggesting dynamic interaction with a restructured microenvironment, potentially supporting matrix stiffening, invasion, and local tumour progression.

Growth factor and mitogenic signalling were active in this subtype, with strong enrichment of MAPK and Ras pathway components (*MAPK3, MAPK10, FGFR2/3, ERBB3, PDGFB*), and a broad panel of upstream ligands and adaptors (*EGF, IGF1, FGF9, FGF18, NTRK2*). These cascades likely reflect growth factor responsiveness and contribute to epithelial maintenance and proliferation.

Metabolic adaptation and insulin signalling also featured prominently. Subtype *Gamma* showed upregulation of genes involved in nucleotide biosynthesis (*NT5C2, GMPR*), redox balance (*IDH1, SUOX*), and amino acid/lipid metabolism (*GLS2, ACACB, GUCY1B1*). Oxidative phosphorylation genes were strongly expressed, including mitochondrial respiratory chain components (*SDHA, COX6A1, MT-ND5, MT-CO1*), suggesting high mitochondrial activity and ATP production. Genes related to insulin signalling and glucose transport (*INSR, SLC2A1, SLC2A4, PIK3R3, PCK1, SOCS3*) were also upregulated, along with enrichment of diabetes-related pathways (e.g., insulin resistance, type II diabetes). These features suggest a metabolic programme tailored for efficient energy use, anabolic biosynthesis, and glucose responsiveness.

Finally, axon guidance signalling was also enriched, driven by expression of semaphorins (*SEMA3B, SEMA4A, SEMA6D*), ephrins and their receptors (*EFNA1, EFNA4, EPHA1, EPHB3*), and netrin–UNC5 components (*NTN1, UNC5A, UNC5B*). While these molecules classically steer neuronal growth cones, their role in epithelial cells is unclear.

In summary, subtype *Gamma* defines a transcriptionally distinct, epithelial-like tumour state, marked by strong cell–cell adhesion, ECM interaction, and metabolic adaptation, distinguishing *Gamma* from the proliferative (*Alpha*) and inflammatory (*Beta*) subtypes.

### Transcriptional subtype Delta—synaptic-like signalling

Rank *k* = 4 NMF stratification subdivided the original *Gamma* cluster, giving rise to a fourth subtype, *Delta* (**Figures 2c** and **S1c**). To characterise the transcriptional basis of this divergence, we performed differential expression analysis between *Delta* OCMs and the remaining *Gamma*-labelled models. In this comparison, genes called as upregulated are specifically enriched in *Delta* relative to *Gamma*, while genes called as downregulated are more highly expressed in the *Gamma*-labelled group (**Figure S2b**). This approach allowed us to identify the molecular features that distinguish the emergent *Delta* cluster from its *Gamma* progenitor.

Pathway enrichment of *Delta*-high genes (**Figure 3a**) revealed a distinctive programme centred on vesicle-mediated communication, with GO terms such as chemical synaptic transmission, trans-synaptic signalling, synaptic vesicle exocytosis, and cell junction (**Table S2**). Upregulated genes included core components of the neuronal secretory apparatus—*SYP*, *STXBP5L*, *RPH3A*, and *CADPS* (vesicle docking and Ca²⁺-triggered release); *ITPKA* and *CACNG7* (intracellular calcium signalling); and synaptic organisers *SEZ6*, *LHFPL4*, and TNR, which mediate adhesion and axon guidance (**Figure 3f**). Supporting this architecture, *RASD2* and *ARHGAP31* suggest plasticity through small-GTPase signalling and cytoskeletal remodelling, while *FHL1* likely contributes junctional scaffolding.

Additional *Delta*-specific genes reinforce this psuedoneuronal identity. *DAB1*, *DRGX*, and *GNAL* are involved in axon guidance and GPCR signalling; *MEIS3* is a neural-crest/homeobox regulator; *TAGLN3* and SPIRE1 modulate actin to support neurite outgrowth; *GDAP1L1* participates in mitochondrial dynamics in neurons; and RCSD1 encodes a brain-expressed cytoskeletal adaptor. CollecTRI analysis also identified RE1-silencing transcription factor (REST) as significantly less active in subtype *Delta* compared with *Gamma* (**Figure S2c**). REST is a well-established transcriptional repressor of neuronal gene expression in non-neuronal tissues. Reduced REST activity in *Delta* OCMs may therefore permit de-repression of neuronal and synaptic genes, contributing to the vesicle-based, pseudoneuronal transcriptional programme that distinguishes this cluster.

In summary, subtype *Delta* appears to represent a transcriptional state whereby neuronal-like secretory machinery is co-opted to support vesicle-based communication, paracrine signalling, or microenvironmental crosstalk. This programme distinguishes *Delta* from its *Gamma* progenitor and highlights a previously underappreciated axis of vesicle-driven signalling and neuronal mimicry within HGSOC.

### Survival analysis

As survival differences have previously been reported for bulk RNA-defined subtypes of HGSOC, we investigated whether similar associations existed in our OCM cohort. We assessed overall survival across the transcriptional subtypes identified at *k* = 2, 3, and 4. Survival was measured from the date of diagnosis to the date of death, and statistical comparisons were performed using Kaplan–Meier analysis with log-rank testing (**Figures 3g** and **S3**). No significant survival differences were observed at any cluster resolution. This suggests that, within this cohort, the transcriptional subtypes captured by NMF do not stratify prognosis based on survival from diagnosis.

### Temporal resolution of transcriptional subtypes revealed by longitudinal OCMs

To determine whether transcriptional subtypes remain stable during disease progression, we analysed longitudinally derived OCMs—models established from the same patient at different timepoints. Because the original NMF clustering was performed using one OCM per patient to avoid patient-specific bias, these additional samples were not part of the transcriptional subtype discovery.

PCA revealed a broad spectrum of within-patient variation (**Figure 4a**). While some patients, e.g. 66, showed tight clustering of their OCMs in PCA space, suggesting transcriptional stability over time, others exhibited marked divergence, with samples occupying distant regions of the PCA plot, consistent with transcriptional divergence during the disease course. For example, for patient 341, OCM.341.1 was generated from a sample collected prior to treatment, while two additional models, OCMs 341.3 and 341.4 were generated from samples collected after several cycles of carboplatin/paclitaxel chemotherapy and just prior to relapse (**Figure 4b**). All three showed transcriptional divergence in PCA space (**Figure 4a**). Similarly, longitudinal samples from patients 288, 327 and 333 (**Figure S4a**) also moved across the PCA landscape.

**Figure 4.**
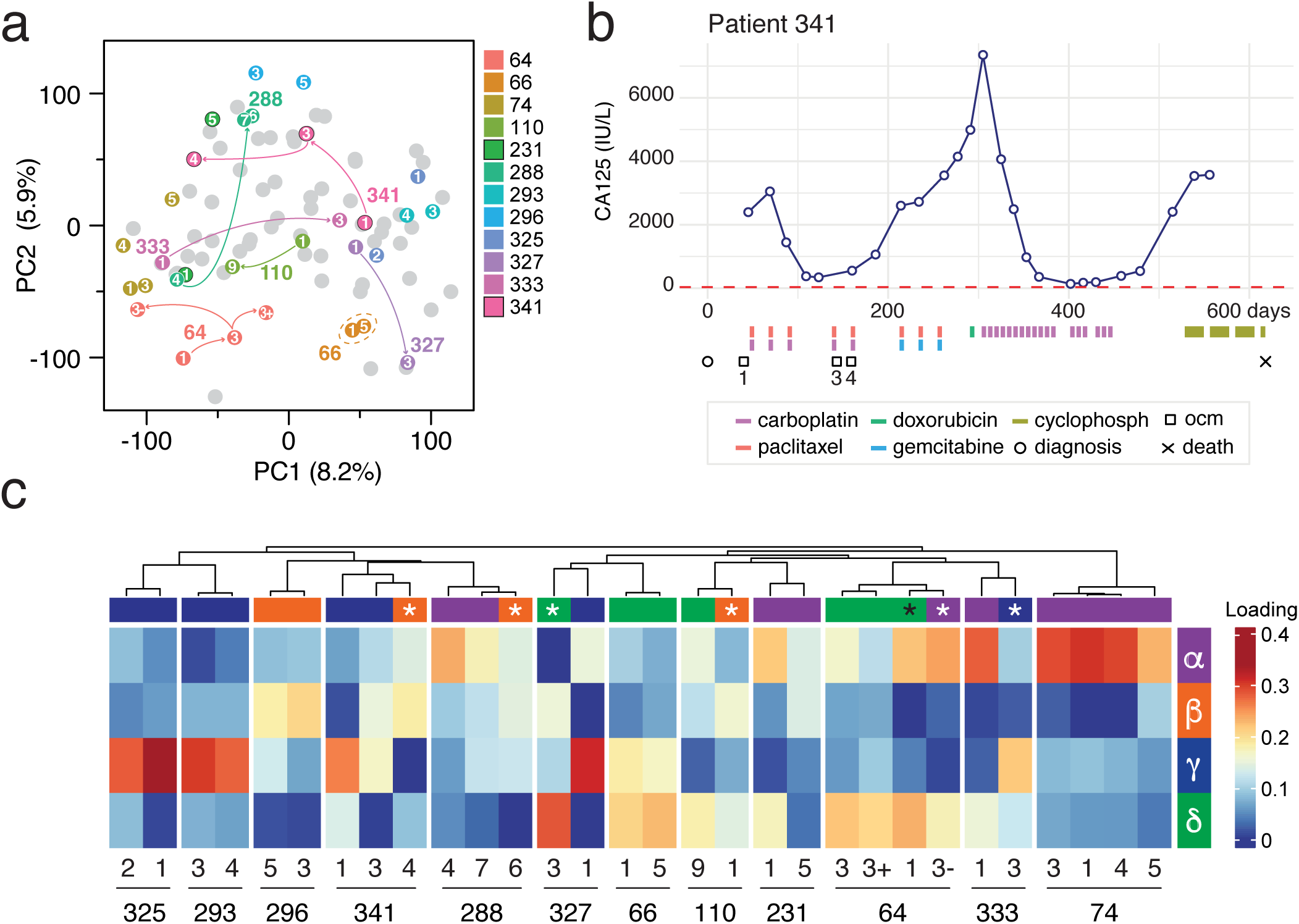
Temporal resolution of transcriptional subtypes reveals intra-patient plasticity. (a) PCA of all 88 tumour-derived OCMs, including longitudinal samples from the same patients. Each colour indicates a distinct patient with ≥2 OCMs; grey dots denote patients with only a single sample. Arrows highlight examples of patient-derived OCMs that shift across PCA space over time, whereas the dashed circle marks patient 66, whose two samples cluster tightly, indicating transcriptional stability. (b) Longitudinal clinical timeline for Patient 341, showing CA125 levels over time (blue line) and annotated with diagnosis (circle), OCM sampling (squares), and death (cross). Coloured bars below indicate timing of chemotherapeutic exposure: purple = carboplatin, red = paclitaxel, bright green = doxorubicin, blue = gemcitabine, and khaki green = cyclophosphamide. (c) Non-negative least squares projection of longitudinal OCM samples into the 4-cluster NMF transcriptional space. Rows represent clusters *Alpha*–*Delta*, and columns represent OCM samples split by patient number and hierarchically clustered. Heatmap colour indicates loading strength (red = high; yellow/white = intermediate; blue = low). Bar above each column denotes the cluster with the highest loading for that sample, with sample ID and sub-sample numbering shown below. White asterisks indicate OCMs that switch transcriptional subtype; black asterisk indicates OCM.64.1, which clustered with *Alpha* by NMF and *Delta* by NNLS.

To determine whether these divergent PCA locations resulted in subtype changes, we projected each longitudinal sample onto the existing NMF cluster space using NNLS (**Figure 4c**). Subtype loading scores were derived using the W matrix from each optimal NMF solution (, = 2, 3 and 4). This allowed us to assess transcriptional subtypes for all longitudinal OCMs, while maintaining the structure of the original clustering.

Several patients (e.g. 66, 293 and 325) showed consistent subtype identity over time, suggesting a stable transcriptional programme. In contrast, others (e.g. 288, 341, 333) exhibited dynamic reconfiguration, with clear shifts in subtype loading across timepoints. For example, OCM’s from patient 327 transitioned from strong *Gamma* to strong *Delta* loading. In other cases, such as 231 and 64, models displayed mixed or transitional subtype identities. OCM.231-5 showed poor loading for all defined clusters, despite chemo-naïve OCM.231-1 showing clear *Alpha* features. This may indicate a novel or less common subtype not captured by the current cluster definitions, suggesting that further granularity may exist beyond the clusters defined in our analysis. Patient 341 exemplifies an interesting and gradual shift in transcriptional subtype. While chemo-naïve OCM.341-1 loaded strongly for *Gamma*, post-treatment OCM.341-4 loaded most strongly for *Beta*, with OCM.341-3 displaying intermediate *Beta* /*Gamma* loadings, consistent with a transition between subtypes or a mixed population.

Together, these results highlight that transcriptional identity is not static within patients. These transitions suggest that subtype plasticity occurs within patients over time, potentially reflecting tumour evolution, treatment-induced selection of subclones, or microenvironmental influence. It also underscores the importance of temporal context in characterising tumour heterogeneity.

### Identification of Cancer Cell Line Encyclopaedia models with HGSOC transcriptional hallmarks

To determine whether the transcriptional subtypes identified above are represented in other collections of ovarian cancer models, we turned our attention to the Cancer Cell Line Encyclopaedia (CCLE). We previously showed that application of NMF to RNA-seq from 44 ovarian cancer cell lines resulted in five clusters that represented the five histological subtypes of epithelial ovarian cancer (Barnes et al., 2021). Classification of these clusters was confirmed by comparison of mutational profiles with clinical cohorts. Collectively, this analysis indicated that of the 44 epithelial ovarian cancer cell lines in total, 16 were likely HGSOC-derived.

First, we updated our classification to include the 24Q4 release of the CCLE, which now includes a panel of 59 epithelial ovarian cancer cell lines. As before, using RNA-seq data from the CCLE (Ghandi et al., 2019), we performed NMF at *k* of 2–10, with 50 NMF runs per,, using the cophenetic correlation coefficient, dispersion coefficient and silhouette width as quality metrics to assess the optimum value of *k* (**Figure S5a**). Again, as per our previous analysis, the five-subgroup model was robust, with the new *k* = 5 clusters aligning well with our original model and with the inferred histological subgroups (**Figure 5**). Of the 44 cell lines analysed previously, 41 remained in the same cluster. EFO27 moved from endometrioid to mucinous, in line with our previous caution that this line shared features of both these subtypes (Barnes et al., 2021). While CAOV3 and FUOV1 are no longer in the HGSOC subgroup, they do cluster with HGSOC in a subset of NMF runs. However, our updated analysis identifies six of the lines new to the 24Q4 release as HGSOC (OV17R, TO14, COV413A, SNU251, UWB1289, PEO1). Thus, we conclude that the CCLE contains 19 cell lines with the transcriptional hallmarks of HGSOC.

**Figure 5.**
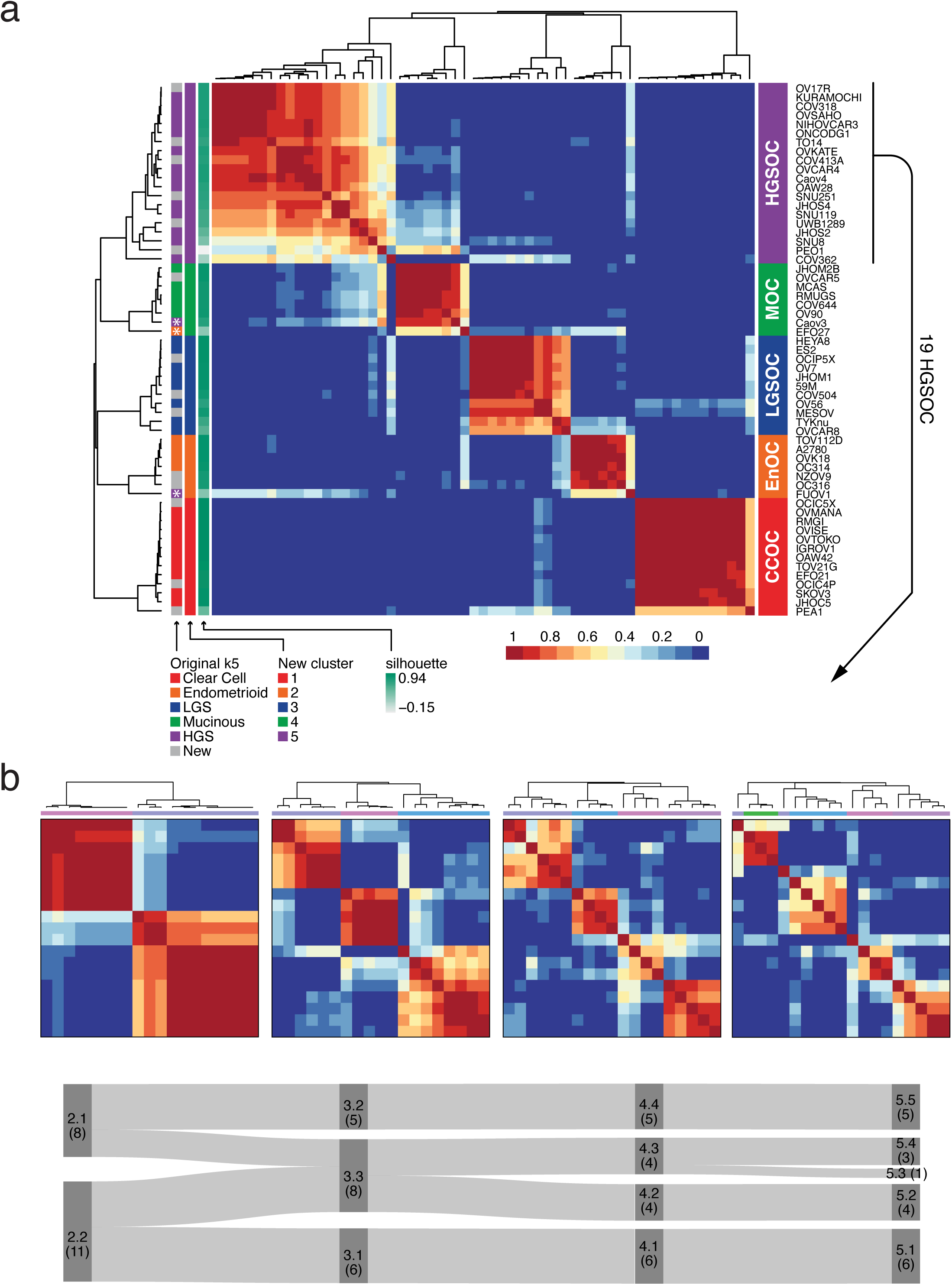
Transcriptional hierarchy is recapitulated in established ovarian cancer cell lines. (a) NMF consensus clustering map of epithelial ovarian cancer cell lines from the Cancer Cell Line Encyclopaedia (CCLE), extending the analysis from Barnes et al., 2021. The consensus heatmap reflects clustering probabilities across NMF runs (, = 5). Annotation bars (from left to right) indicate: original study cohort membership (coloured vs grey for new), cluster assignment from this analysis (, = 5) and silhouette score. To the right, annotation bar represents inferred histopathological subtype for each line. HGSOC-classified lines (n = 20) are indicated by line. ONCODG1 is later removed as it has been identified to be a derivative of OVCAR-3 cell line (Capes-Davis et al., 2010), leaving n =19 HGSOC cell lines. (b) NMF clustering of the 19 CCLE HGSOC cell lines. Consensus matrices for *k* = 2 to *k* = 5 are shown, based on repeated runs using 11 variable gene lists. Heatmaps reflect clustering consistency across 200 NMF iterations, with red = high consensus and blue = low. (c) Sankey diagram tracing cluster membership across different number of clustering resolutions. Cell line cluster assignment was based on the most frequent (modal) cluster membership observed across 11 NMF runs using different sizes of input gene list based on median absolute deviation.

### Transcriptional hierarchy is recapitulated in established ovarian cancer cell lines

With the expanded collection of 19 HGSOC cell lines, we next asked whether the transcriptional hierarchy observed in patient-derived OCMs is recapitulated in established HGSOC cell-line models. The 19 HGSOC lines were subjected to the same NMF pipeline: 11 input matrices (500–10,000 most-variable genes) were analysed at ranks *k* = 2–10 using 50 iterations, and quality metrics were compared. Cophenetic, silhouette and dispersion scores (**Figure S6a**) again pointed to a robust two-cluster backbone, with variable indications of stability across k = 3 to 5. We repeated NMF for *k* = 2, 3, 4 and 5 using 200 iterations and all 11 input gene lists. Inspection of cluster membership across all gene lists showed that cell-line assignments were highly consistent (**Figure S6c**). A Sankey plot summarising cell line flows confirmed a hierarchical pattern: some cell lines from the two parental clusters first merge to create a third mixed cluster at *k* = 3 (3.3), mirroring the *Alpha* /*Beta* /*Gamma* configuration in the OCM biobank; at *k* = 4 this mixed cluster resolves cleanly into two daughter clusters (4.3 and 4.2) whose membership traces back to each of the original *k* = 2 clusters ; and at *k* = 5 a single cell line splits off to form a small fifth group (**Figure 5c**). Crucially, no cell line ever switches idiosyncratically or becomes a singleton before the final split, underlining the stability of the hierarchy across increasing ranks.

Thus, the HGSOC cell-line panel also exhibits a nested transcriptional architecture as seen in OCMs: an initial two-cluster scaffold, three clusters formed by some merging of those scaffolds, and orderly splits at higher ranks. This consistency—despite the relatively small panel of lines—strengthens the biological credibility of the inferred transcriptional hierarchy seen in the OCM cohort.

## DISCUSSION

In this study, we leveraged a living biobank of highly purified patient-derived OCMs to revisit the long-standing question of whether HGSOC harbours stable, tumour-cell-intrinsic transcriptional subtypes. Prior work using bulk tumour profiling described four molecular subtypes (Tothill et al., 2008; TCGA, 2011). However, subsequent single-cell and spatial transcriptomic studies have suggested that only two of these, the C5/Proliferative and C4/Differentiated groups, likely reflect tumour-cell-intrinsic states; the remaining subtypes, C2/Immunoreactive and C1/Mesenchymal, appear to be strongly confounded by infiltrating immune and stromal cells (Izar et al., 2020; Olbrecht et al., 2021; Deng et al., 2022).

By applying NMF to RNA-seq profiles from highly-purified patient-derived tumour-cell fractions, we identified robust transcriptional subtypes without the confounding effects of stromal admixture. Across a range of input gene sets and clustering resolutions (, = 2–4), our results revealed a stable, hierarchical architecture that goes beyond this dichotomy. At *k* = 2, models separated cleanly into two groups; at *k* = 3, a third cluster emerged through reorganisation of samples from both parent clusters, indicating transcriptional heterogeneity not captured at lower resolution. Intriguingly, at *k* = 4, we observed a consistent and hierarchical split of ***Gamma*** into two clusters. This gave rise to a fourth cluster, ***Delta***, that is not simply an artefact of overclustering as it emerged reproducibly across gene inputs and did not disrupt the overall structure of the hierarchy. In contrast to the ***Beta*** cluster, which emerged via recombination, ***Delta*** branched cleanly from ***Gamma***, suggesting it represents a nested, transcriptionally distinct state within a broader epithelial-like lineage.

Taken together, these findings suggest that HGSOC is characterised by three core tumour-cell-intrinsic transcriptional subtypes. The ***Alpha*** cluster was enriched for cell-cycle and DNA replication machinery, strongly aligning with the C5/Proliferative signature described in bulk tumour studies. While the ***Beta*** cluster exhibits immune-related features, it is distinct from the immune-cell-infiltrated C2/Immunoreactive subtype described in prior bulk sequencing datasets. Specifically, ***Beta*** demonstrated a tumour-cell-intrinsic inflammatory programme, defined by cytokine signalling, NF-κB activation, and immune mimicry. ***Gamma*** exhibited markers of epithelial differentiation, including classical lineage markers and metabolic signatures, echoing features of the C4/differentiated subtype—with an additional finer-grained programme, ***Delta***, nested within. This hierarchical structure refines existing subtype models and underscores the potential of using purified tumour models to uncover tumour-cell-intrinsic transcriptional diversity in ovarian cancer.

The ***Alpha*** cluster, while aligning with the previously described C5/Proliferative subtype, exhibits a more coherent and intensified transcriptional programme. We observed coordinated upregulation across the full cell-cycle axis—G1/S transition, DNA replication and mitosis—alongside elevated expression and inferred activity of E2F1–4. E2F activity is usually repressed by the DREAM complex, composed of Rb-family proteins (Rb, p107, p130), E2F4/DP, and the MuvB complex. In response to mitogenic signals, Cyclin D–CDK4/6 and Cyclin E–CDK2 phosphorylate Rb and DREAM components, disassembling the complex and releasing E2F to drive S-phase entry genes such as MYBL2 (B-Myb). The MuvB complex then transitions to an activating role through sequential interaction with B-Myb and FoxM1, coordinating the expression of genes required for mitosis and late cell-cycle progression (Iness and Litovchick, 2018). In HGSOC, elevated *MYBL2* expression is significantly associated with advanced-stage disease and poorer overall survival (Iness et al., 2021). Similar “E2F-high” states have been described in other malignancies: *MYBL2* is incorporated into breast cancer recurrence-risk signatures (Solin et al., 2013), and E2F target gene enrichment identifies a poor-outcome subtype in pancreatic ductal adenocarcinoma (Lan et al., 2018).

In ***Alpha*** OCMs, the transcriptional intensity of this programme suggests that this axis may be constitutively active or uncoupled from upstream checkpoint control. While we did not assess genomic alterations directly, the expression profile is congruent with known features of HGSOC—such as frequent Rb pathway dysfunction and *CCNE1* amplification. Taken together, these findings position the ***Alpha*** subtype as a transcriptionally stressed, checkpoint-compromised phenotype. This raises the hypothesis that ***Alpha*** tumours could be selectively vulnerable to agents targeting the cell cycle or mitotic stress response, such as CDK inhibitors, CHK1/WEE1 blockade, or PLK1 inhibition.

Notably, ***Alpha*** models also displayed hallmarks of EMT, suggesting possible acquisition of invasive traits alongside proliferative deregulation. This dual phenotype — of unchecked proliferation and emerging plasticity — may represent a particularly aggressive tumour cell state, potentially associated with rapid progression or early relapse. Whether cell cycle deregulation and EMT coevolve independently, concurrently or whether one is facilitated by the other remains unclear.

The chronic inflammatory programme displayed by ***Beta*** is characterised by IL1/IL6 cytokines and full engagement of the MyD88–TRAF–NF-κB axis. Chronic inflammation is increasingly recognised as a driver of tumour progression in HGSOC (Savant et al., 2018), with pro-inflammatory cytokines such as IL-6 and TNF-*Alpha* frequently upregulated and shown to promote growth, metastasis, and treatment resistance (Pawlik et al., 2021). These signals converge on NF-κB and STAT3 pathways, which enhance cell survival, proliferation, and resistance to apoptosis (Tosic and Frank, 2021; Devanaboyina et al., 2022). Clinically, overactive NF-κB has been linked to reduced platinum sensitivity and shorter progression-free survival (Kan et al., 2020), while preclinical inhibition of NF-κB or related JAK/STAT3 signalling can re-sensitise tumours to therapy (Wu et al., 2019).

Beyond intrinsic tumour signalling, NF-κB drives immune modulation, regulating expression of cytokines and chemokines that recruit tumour-associated macrophages (TAMs) and regulatory T cells (Tregs), both of which suppress cytotoxic immunity (Liu et al., 2025). NF-κB also induces immune checkpoints such as PD-L1, which suppresses T cell activity via PD-1 engagement. In the *Beta* cluster, PD-L2—the alternative ligand for PD-1—is significantly upregulated, pointing to a similar immune-evasive phenotype. PD-L2 has recently emerged as a prognostic marker and potential immunotherapy target in HGSOC (Yates et al., 2024), and PD-1/PD-L1 inhibitors are under clinical evaluation (Chen et al., 2024). Together, these features position NF-κB as a central node linking inflammation, immune mimicry, and immunosuppression in the ***Beta*** subtype.

The ***Gamma*** cluster displayed a well-differentiated epithelial phenotype, marked by high expression of canonical lineage markers including *PAX8*, *WT1*, *EPCAM*, and *MUC16* (CA125). This expression pattern is consistent with the C4/Differentiated subtype previously described in bulk tumour studies, and aligns with transcriptional profiles of secretory fallopian tube epithelium (Tothill et al., 2008; Verhaak et al., 2013; Lengyel et al., 2022). CA125/MUC16 expression was particularly elevated, suggesting that this cluster may correspond to the high-CA125 phenotype frequently observed in HGSOC (Tothill et al., 2008; Verhaak et al., 2013). Retention of epithelial features, including strong E-cadherin expression and suppression of mesenchymal markers such as vimentin, further underscores the non-EMT-like character of this group. Interestingly, these same traits have been linked to more favourable clinical outcomes in HGSOC. For example, low vimentin expression and preserved epithelial architecture have both been associated with longer overall survival in retrospective analyses (Konecny et al., 2014; Gonzalez et al., 2018). Taken together, our data support the existence of an epithelial–metabolic subtype within HGSOC, characterised by CA125 overexpression, epithelial lineage fidelity, and oxidative metabolism. This state is transcriptionally distinct from the proliferative and inflammatory programmes described above, and may confer therapeutic vulnerabilities (e.g. metabolic dependencies)

The ***Delta*** subtype up-regulates synaptic vesicle, axon-guidance, and Ca²⁺-cycling genes (*SYP*, *STXBP5L*, *SEZ6*, *CACNG7*), implying that some tumour cells co-opt neuronal-like secretory programmes to modulate their microenvironment through vesicle-based communication. This echoes recent findings that platinum stress can unlock a neuronal/stem-like axis in ovarian cancer cells (Diaz-Carballo et al., 2024). Important next steps will be functional interrogation of OCMs from this subtype to compare synaptic vesicle biology and secretory capacity with the other transcriptional subtypes.

By projecting longitudinal OCMs into the subtype framework, we observed both clonal stability and flexible reprogramming. Some tumours retained a fixed state, suggesting dominant clones that persist despite treatment, while others shifted from differentiated-like ***Gamma*** to proliferative ***Alpha*** or immune-mimicry ***Beta*** states. This raises the possibility that the transcriptional subtypes do not reflect disease subtypes defined during the early stages of HGSOC evolution. Indeed, these transcriptional shifts raise key questions: Are subtypes rewired by the microenvironment or treatment, or do pre-existing clones expand under selective pressure — or both? Future work combining single-cell lineage tracing, co-cultures, epigenetic profiling and *in vitro* evolution could reveal whether these transitions are hardwired or reversible.

While our study provides a refined view of tumour-cell-intrinsic transcriptional subtypes in HGSOC, several limitations should be acknowledged. Although the use of highly purified, patient-derived OCMs enables interrogation of tumour-cell–intrinsic programmes without stromal confounding, by definition it excludes the complex microenvironmental interactions—such as tumour–stroma crosstalk—that shape tumour behaviour *in vivo* and are not recapitulated in monoculture. Furthermore, despite being low-passage and retaining strong transcriptional resemblance to the originating tumours, OCMs may still undergo selective pressures *in vitro* that favour certain subpopulations, potentially introducing transcriptional biases. The number of models per subtype—particularly ***Delta***—remains modest, limiting power to draw definitive conclusions about subtype-specific clinical associations or therapeutic vulnerabilities. Finally, while we propose candidate vulnerabilities, such as cell cycle checkpoint dependence in ***Alpha*** or immunomodulatory features in ***Beta***, functional validation will be needed to determine whether these features translate into actionable therapeutic opportunities.

Taken together, this study delivers the first cohort-scale, tumour-cell-intrinsic roadmap for transcriptional heterogeneity in HGSOC—free from stromal noise and microenvironmental confounders. This refined framework establishes a new benchmark for decoding the hidden complexity of HGSOC and offers a clear path forward for functional validation, precision patient stratification, and the rational design of subtype-specific therapies.

## MATERIALS AND METHODS

### Contact for reagent and resource sharing

Further information and reagent requests may be directed to Stephen S. Taylor.

### Patient samples

Research samples were obtained from the Manchester Cancer Research Centre (MCRC) Biobank, UK. The MCRC Biobank holds a generic ethics approval which can confer this approval to users of banked samples via the MCRC Biobank Access Policy. Further details regarding ethical governance and licensing can be found at: https://www.mcrc.manchester.ac.uk/research/mcrc-biobank/accessing-the-mcrc-biobank/.

### Establishment and culture of ovarian cancer models

Establishment of patient-derived OCMs and matched stromal fibroblasts has been described previously (Ince et al., 2015; Nelson et al., 2020). For ascites, samples were centrifuged at 500 × g for 10 minutes at 4 °C, and the resulting cell pellets were pooled in HBSS (Life Technologies). Red blood cells were lysed using a commercial lysis buffer (Miltenyi Biotec) according to the manufacturer’s instructions. The remaining cells were seeded into Primaria flasks (Corning) containing OCMI medium (Ince et al., 2015). Solid tumour specimens were enzymatically dissociated using the Miltenyi Tumour Dissociation Kit, again following the manufacturer’s protocol. Dissociated cells were plated into collagen-coated 12.5 cm² flasks containing OCMI. Cultures were maintained at 37 °C in a humidified atmosphere with 5% CO₂ and 5% O₂ for 2–4 days. Media changes were performed every 3–4 days. Once tumour cell adhesion was established, stromal cells were selectively detached using 0.05% trypsin-EDTA and transferred to gelatin-coated flasks in OCMI supplemented with 5% FBS. Tumour cells were cultured until reaching ∼95% confluency, at which point they were passaged using 0.25% trypsin-EDTA, centrifuged in DMEM with 20% FBS, and re-plated at a 1:2 ratio and maintained in OCMI medium. For long-term preservation, cells were cryopreserved in Bambanker (Wako Pure Chemical).

### OCM *TP53* transcript sequencing

Total RNA was extracted using the RNeasy Plus Mini Kit (Qiagen), and *TP53* cDNA was synthesised using the SuperScript™ III One-Step RT-PCR System with Platinum Taq HiFi (Thermo Fisher Scientific) and the following primers: forward 5’-CAC CTC GAG GAG GAG CCG CAG TCA GAT CCT A-3’ and reverse 5’-CAC GCG GC CGC TCA CAG TCT GAG TCA GGC CCT TCT GTC-3’. Amplified PCR products were cloned into the pBluescript SK-vector and transformed into XL1-Blue competent cells. Plasmid DNA was purified using the QIAprep Spin Miniprep Kit (Qiagen) and sequenced using the following primers: 5’-CAC CAG CAG CTC CTA CAC CG-3’, 5’-ATG AGC GCT GCT CAG ATA GCG-3’, 5’-CGG CTC ATA GGG CAC CAC C-3’, 5’-TCT TCT TTG GCT GGG GAGAGG-3’. Sequence alignments were performed using SeqMan Pro (DNASTAR). For newly derived OCMs described in this study, pBluescript-p53 plasmids were subjected to full-length sequencing via Oxford Nanopore (Plasmidsaurus).

### Amplicon-EZ sequencing

Targeted sequencing of *TP53* exon 7 was performed using Amplicon-EZ next-generation sequencing (Azenta Life Sciences). Genomic DNA was extracted from OCMs using the PureLink™ Genomic DNA Mini Kit (Thermo Fisher Scientific), followed by nested PCR amplification using exon 7-specific primers carrying partial Illumina® adapter overhangs. PCR was conducted using RedTaq® DNA polymerase on a ProFlex™ PCR System (Thermo Fisher Scientific). Outer forward primer: 5’-CAG GGG TCA GAG GCA AGC AGAG-3’; outer reverse primer: 5’ TGC CAC AGG TCT CCC CAA GG-3’; inner forward primer with partial adapter overhang: 5’-ACA CTC TTT CCC TAC ACG ACG CTC TTC CGA TCT GGG TGG CAA GTG GCT CCT GAC-3’; inner reverse primer with partial adapter overhang: 5’-GAC TGG AGT TCA GAC GTG TGC TCT TCC GAT CTG CGC ACT GGC CTC ATC TTG G-3’.

Amplicons were assessed for correct size on a 3% 3:1 NuSieve agarose gel, visualised via UV transillumination (UVP), and subsequently purified using the QIAquick PCR Purification Kit (QIAGEN). A total of 500 ng of purified PCR product in EB buffer was submitted for Amplicon-EZ library preparation and sequenced on an Illumina® NovaSeq X platform at Azenta’s European Genomics Headquarters (Leipzig, Germany). For stromal:tumour titration experiments, stromal and tumour derivatives of OCM.231-1 were cultured separately as described above. Once confluent, cells were trypsinised, counted, and mixed at defined ratios: 1:99, 2.5:97.5, 5:95, 60:40, 80:20, 90:10, and 100% stromal and 100% tumour. Mixed cell suspensions were sequenced as above.

### RNA sequencing of ovarian cancer models

RNA-seq of our earlier OCMs has been published (Nelson et al., 2020; Barnes et al., 2021; Coulson-Gilmer et al., 2021; Littler et al., 2025; Tighe et al., 2025). For RNA-seq of OCMs first described here, total RNA extracted using a RNeasy Plus Mini kit (Qiagen) was submitted to the Genomic Technologies Core Facility at the University of Manchester. After quality and integrity assessment using a 4200 TapeStation (Agilent Technologies), libraries were generated using the Illumina® Stranded mRNA Prep Ligation kit (Illumina, Inc.) according to the manufacturer’s protocol. Following a single adenine base addition, adapters with a complementary thymine overhang were ligated to the cDNA fragments followed by ligation of pre-index anchors to prepare for dual indexing. Index adapter sequences were added by PCR to enable multiplexing of the final cDNA libraries, which were pooled and loaded onto an SP flow cell and paired-end sequenced (59 + 59 cycles, plus indices) on an Illumina NovaSeq6000 instrument. Raw sequencing output was demultiplexed (allowing one mismatch), and binary base call (BCL)-to-Fastq conversion performed using bcl2fastq software (Illumina, Inc., v2.20.0.422).

### Data processing of ovarian cancer model RNA sequencing

Adapter sequences and low-quality bases were trimmed using BBDuk from the BBMap suite (v36.32) (Sourceforge brian-jgi, 2025). Cleaned reads were aligned to the human reference genome (hg38) and with gene annotation from Gencode (v32) using STAR (v2.7.2b) (Dobin et al., 2013). Per-gene read counts were generated using the --quantMode GeneCounts parameter within the STAR alignment step.

### Acquisition and pre-processing of Cancer Cell Line Encyclopaedia RNA sequencing

An RNA-seq raw count matrix (22Q4 release) was obtained from the DepMap portal (https://depmap.org/portal). To correct for batch effects associated with differences in library preparation methods (stranded vs. unstranded), we applied ComBat-seq, a normalisation algorithm designed for batch correction of count-based RNA-seq data (Zhang et al., 2020). Cell lines were then filtered to those identified by literature search as epithelial ovarian cancer in origin. The resulting expression matrix was used for downstream analysis.

### Non-negative matrix factorisation of ovarian cancer models

Gene count matrices were transformed using the variance stabilising transformation (VST) from the DESeq2 package (v1.26.0) (Love et al., 2014). To assess the influence of input gene selection on clustering outcomes, we generated up to 11 input matrices, each comprising the top 500 to 10,000 most variable genes (in 1,000-gene increments) as determined by median absolute deviation (MAD). For each gene set, NMF was performed using the NMF R package (v0.23.0) (Gaujoux and Seoighe, 2010) with rank (,) ranging from 2 to 10. Each NMF run was initially performed with 50 iterations using random initialisation points. Clustering performance was assessed using established metrics, including the cophenetic correlation coefficient, average silhouette width, and the dispersion coefficient, calculated across the consensus matrix for each gene list and rank (,). For select ranks, we performed an extended NMF analysis comprising 200 iterations per each of the input gene matrices.

To align clusters across different gene sets (accounting for label switching between runs), we implemented a custom matching strategy using normalised mutual information (NMI). NMI scores were calculated between each pair of clusters using the aricode R package (v1.0.0) (Chiquet, 2024), and a greedy matching algorithm was used to assign consistent cluster identities across runs (**Figures S1c, S5b, S6c**). Final cluster assignments used in downstream analyses were derived from the dominant cluster label for each sample across gene sets.

### Differential gene expression analysis

Differential gene expression analysis was performed using the limma-voom pipeline (Law et al., 2014) to identify genes enriched within each transcriptional subtype. To account for uncertainty in cluster assignment—arising from variation across input gene lists—we calculated the proportion of NMF runs in which each OCM was assigned to its modal cluster. These proportions were then used as observation-level weights in the linear model. For each cluster, genes were tested for differential expression versus the combination of the other clusters. Genes with a Benjamini-Hochberg false discovery rate (FDR) < 0.05 were considered significantly upregulated. The resulting gene lists were used for functional enrichment analysis as described below.

### Pathway enrichment

Pathway enrichment analysis was performed using the g:Profiler2 R package (version 0.2.3) (Kolberg et al., 2023) to functionally characterise genes upregulated in each transcriptional subtype. Differentially expressed gene lists (FDR < 0.05) from each subtype comparison were used as input. Enrichment was assessed across multiple databases, including Gene Ontology (GO) Biological Process, KEGG, and Reactome. The g:SCS algorithm was used to adjust for multiple testing and only terms with an adjusted p-value < 0.05 were retained. Enriched pathways were manually grouped into broader functional themes by literature search to facilitate biological interpretation and visualisation.

### Network analysis

Network analysis was performed in Cytoscape (version 3.10.3) to visualise the interactions between genes enriched in each transcriptional cluster.

### Transcription factor activity

To infer transcription factor (TF) activity from differential gene expression data, we used the decoupleR R package (v2.10.0) and implemented the Univariate Linear Model (ULM) method. Input data consisted of moderated t-statistics from limma-voom differential expression analysis. TF–target interaction data were derived from the CollecTRI prior knowledge network (Muller-Dott et al., 2023). Only TFs with a minimum of five expressed target genes in the dataset were retained. ULM computes TF activity scores by fitting a linear model between each TF’s signed target set and the gene-level test statistics. For each comparison, we retained the top 25 TFs by absolute score.

### Survival analysis

To evaluate the association between transcriptional subtypes and overall survival, we performed Kaplan–Meier survival analysis using the survival and survminer R packages. For each NMF clustering solution (e.g., at rank 2, 3, or 4), the dominant cluster label per sample was treated as a categorical variable. Survival time was defined as days from diagnosis to death or last follow-up. Censoring status was encoded as a binary event variable (1 = death, 0 = censored). We constructed Surv objects using the Surv() function and stratified survival curves by cluster assignment using the survfit() function. Curves were visualised using ggsurvplot(), which included the log-rank p-value and a risk table.

## DATA AVAILABILITY

The published gene expression data presented here have been deposited with EMBL-EBI, accession numbers: E-MTAB-7223; E-MTAB-10801; E-MTAB-11000; E-MTAB-14568. Unpublished gene expression data will be made available upon publication.

## AUTHOR CONTRIBUTION STATEMENT

Barnes: conceptualisation, investigation, formal analysis, methodology, software, visualisation, supervision and writing–original draft. Littler, Tighe, Evans, Altringham, Nelson, Lin and Morgan: methodology, investigation, formal analysis, and resources. McGrail: writing–review and editing and project administration. Taylor: conceptualisation, funding-acquisition, supervision, validation, visualization, writing–review & editing.

## DECLARATIONS

The authors have no competing interests to declare that are relevant to the content of this article.

## ACKNOWLEDGEMENTS

Research in the Taylor lab is funded by Cancer Research UK [C1422/A31334] and [C147/A25254]; the Medical Research Council [MR/X008088/1]; and the NIHR Manchester Biomedical Research Centre [NIHR203308]; the views expressed are those of the author(s) and not necessarily those of the NIHR or the Department of Health and Social Care. We thank the patients for their commitment to research; the MCRC Biobank for the sample collection; members of the Taylor lab for advice and comments on the manuscript.

## SUPPLEMENTARY INFORMATION

**Table S1.**
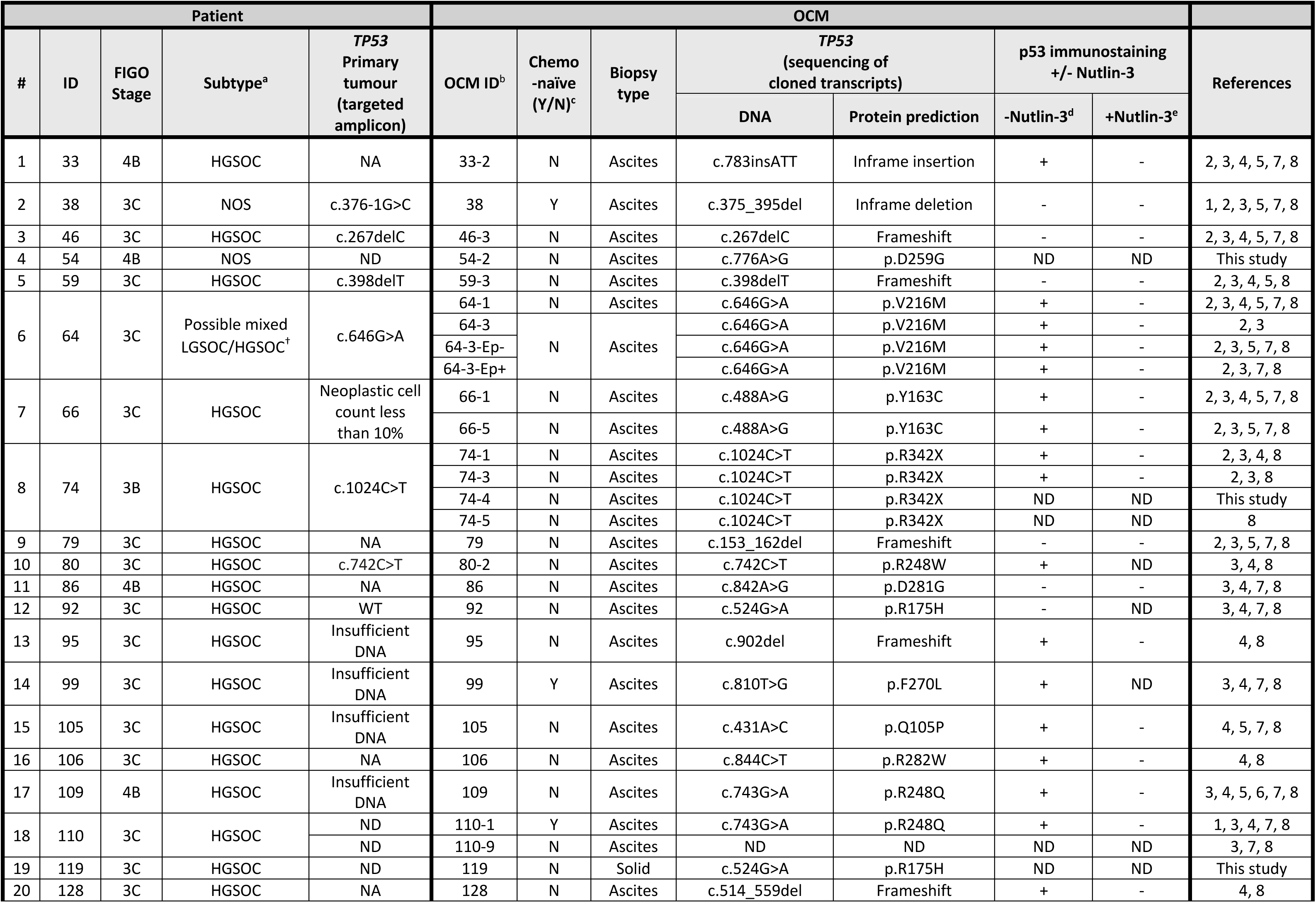
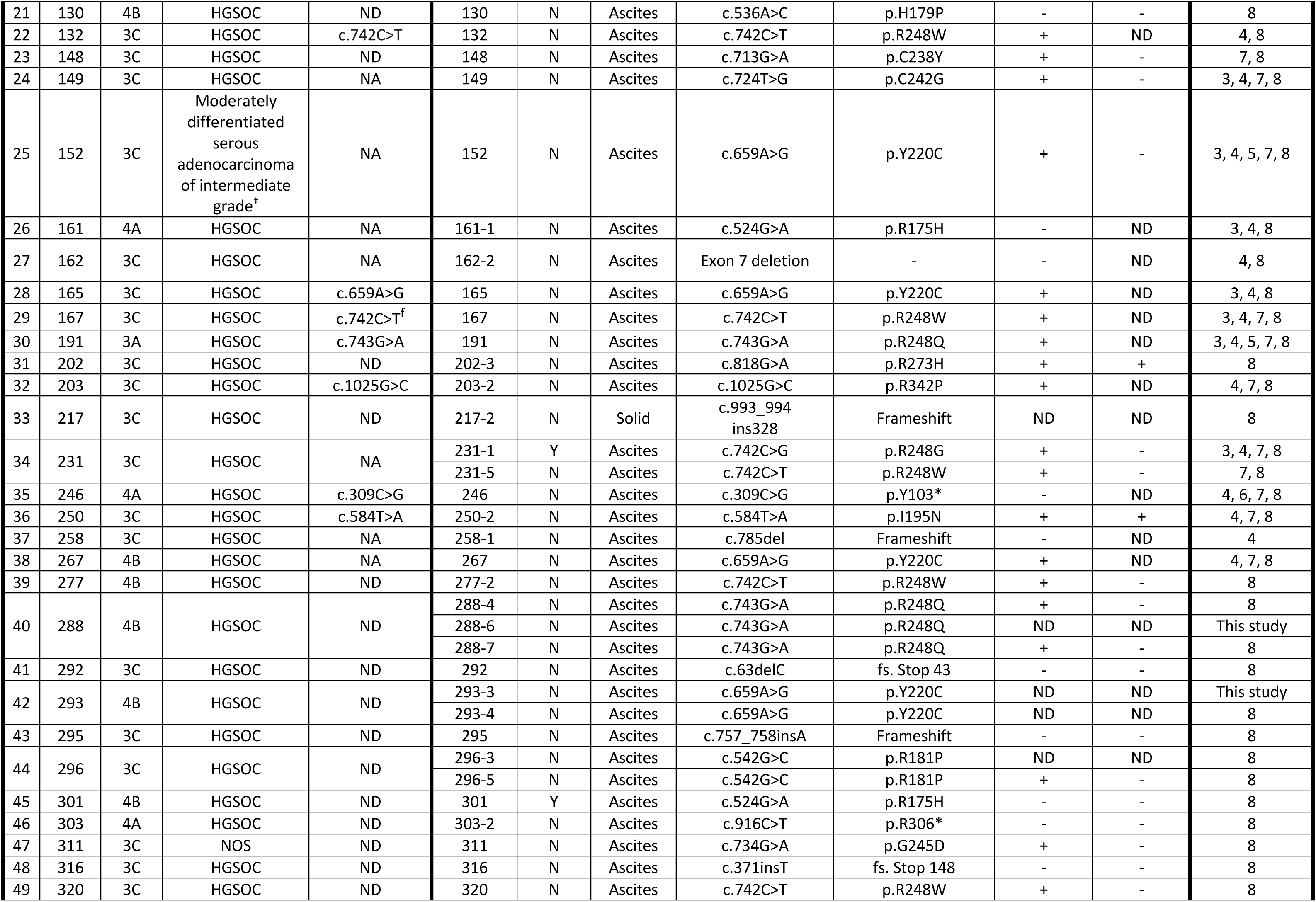
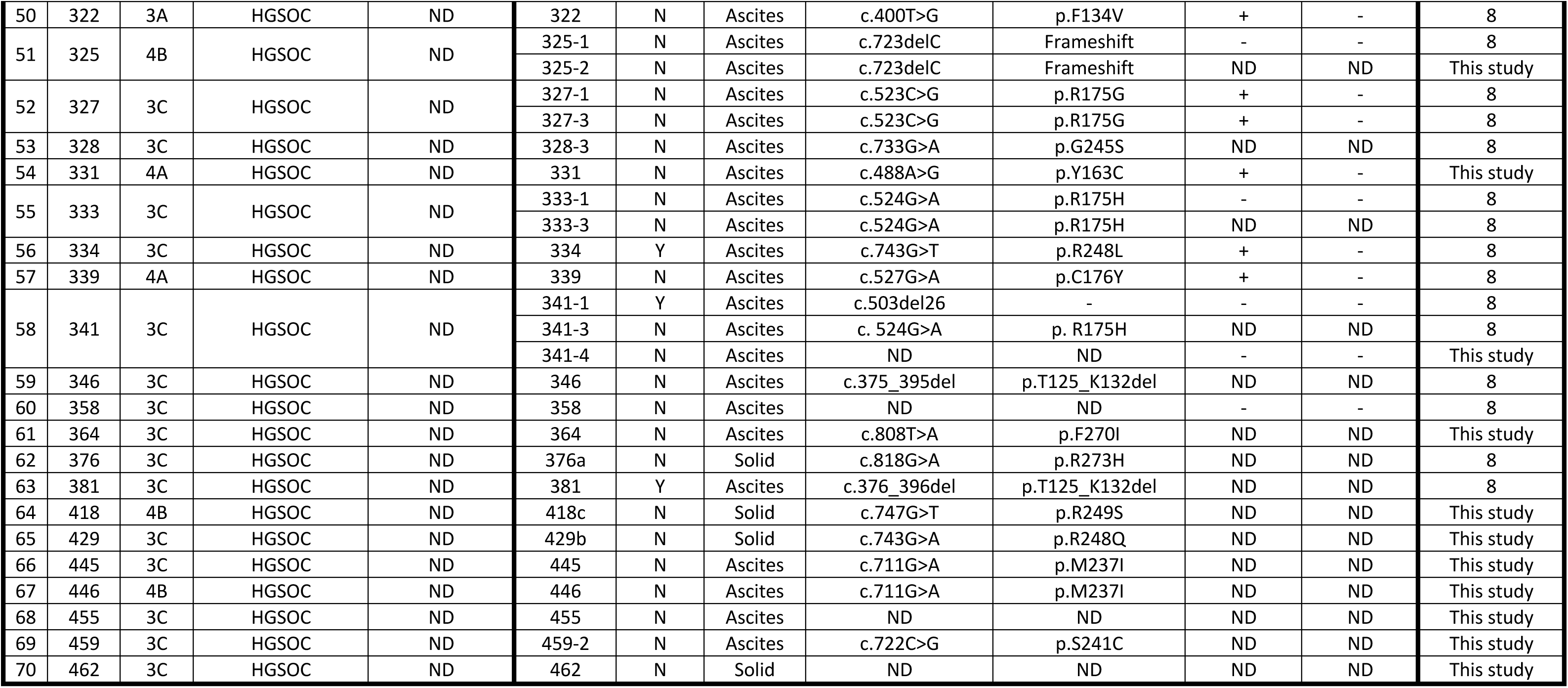
Patient and OCM characteristics.

Table S2. Subgroup pathway analysis

## FIGURE LEGENDS

**Figure S1.**
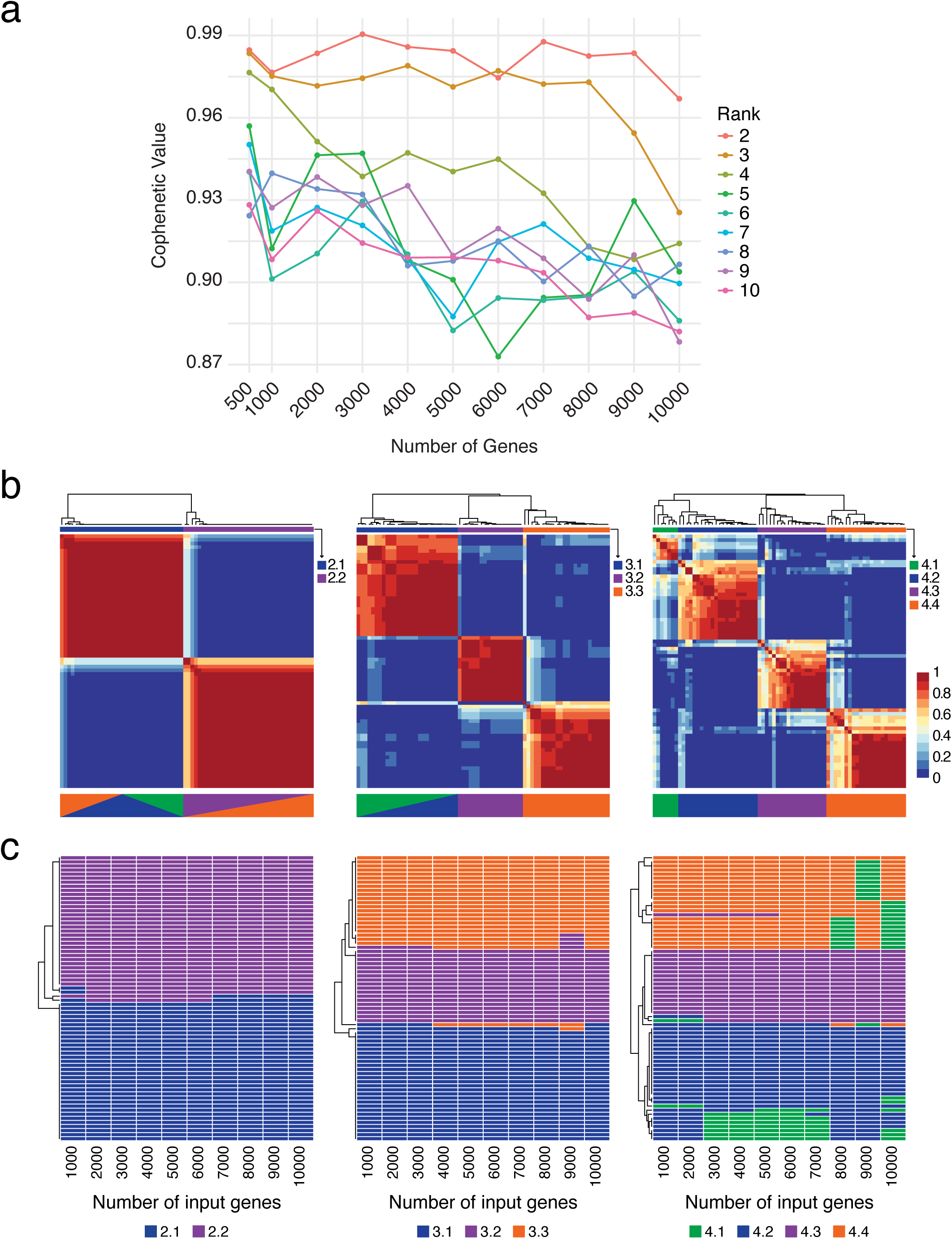
Assessment of clustering quality and consistency. (a) Cophenetic correlation coefficient across input gene list sizes (1,000 to 10,000 genes) and NMF ranks (, = 2 to 10). Each line represents one rank. Cophenetic values were calculated for 50 NMF iterations per condition and used to evaluate clustering stability. (b) Representative consensus matrices for NMF clustering at *k* = 2 (left), *k* = 3 (middle), and *k* = 4 (right) using the 3,000, 6,000, and 4,000 gene input sets, respectively. Consensus clustering maps show pairwise sample co-clustering frequency across 200 NMF runs. Red indicates samples that consistently co-clustered; blue indicates samples that never co-clustered. Cluster assignments and silhouette scores are shown as annotation bars. Below each map indicates the colouring based on the Sankey plot in **Figure 2c**. (c) Sample-wise cluster assignments across gene list sizes for *k* = 2 (left), *k* = 3 (middle), and *k* = 4 (right). Rows represent individual OCMs (n = 70); columns represent input gene lists (1,000 to 10,000 genes). Colours indicate assigned cluster identity. Panels illustrate the consistency of cluster assignment across input sizes, with the emergence of subclusters (e.g. *Delta* at *k* = 4).

**Figure S2.**
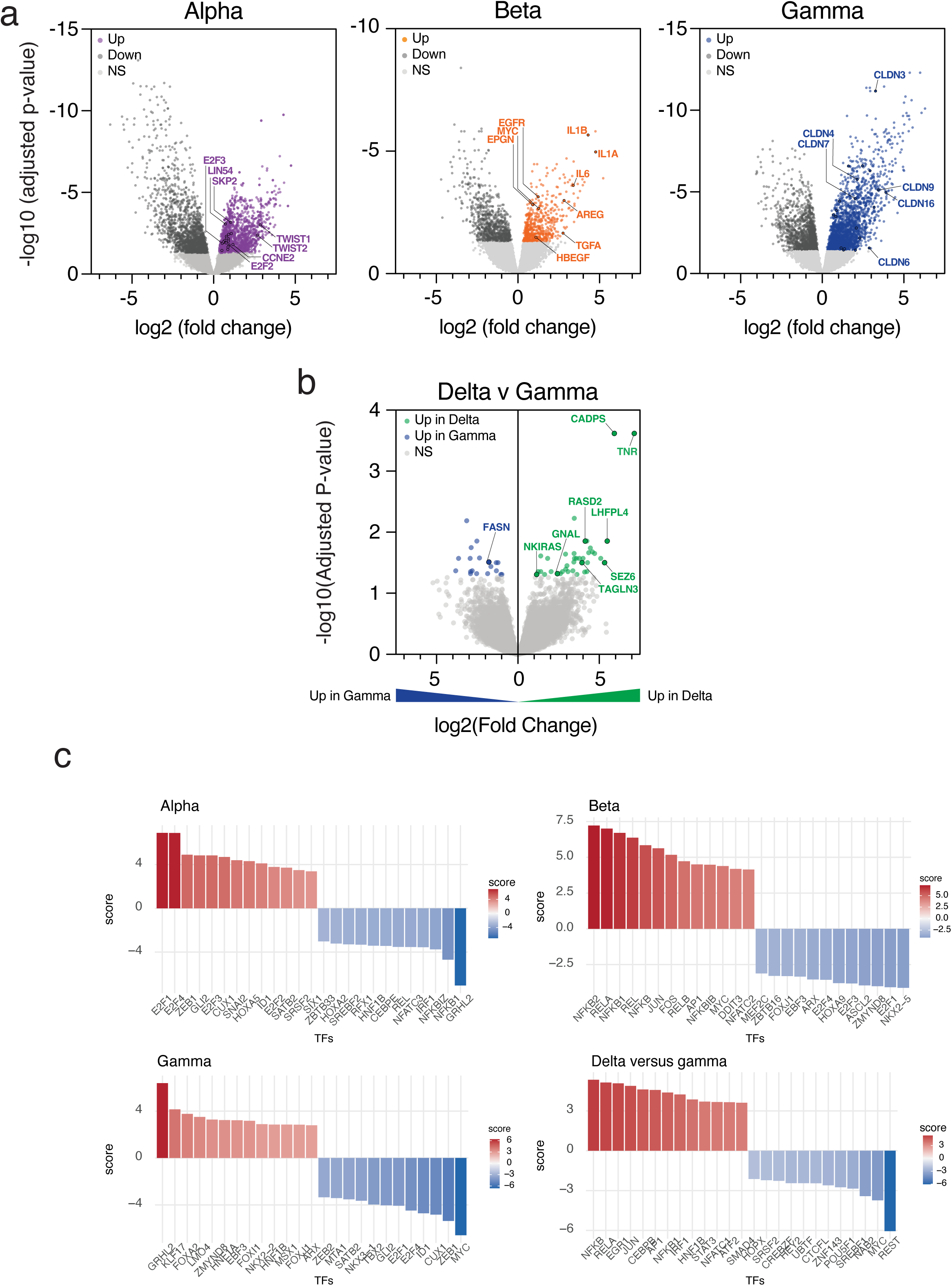
Differential gene expression of transcriptional subtypes. (a) Volcano plots showing log₂ fold change (x-axis) versus –log₁₀ adjusted p-value (y-axis) for genes differentially expressed in each cluster relative to the other two (*Alpha*, left; *Beta*, centre; *Gamma*, right). Differential expression was assessed using limma-voom with sample weights derived from NMF cluster assignment consistency across input gene sets. Genes highlighted in orange, purple and blue are upregulated, dark grey are downregulated, and light grey are not significantly different (FDR ≥ 0.05). (b) Volcano plot comparing subtype *Delta* to *Gamma* from the 4-cluster NMF solution. Genes on the right (green) are significantly upregulated in *Delta*, while genes on the left (blue) are significantly upregulated in *Gamma*. Comparison was performed using limma-voom with the same sample-weighting strategy. Genes not significantly differentially expressed (FDR ≥ 0.05) are shown in grey. Selected genes of interest are labelled based on literature relevance and agreement with downstream pathway enrichment analyses. (c) Transcription factor (TF) activity profiles for *Alpha*, *Beta*, *Gamma*, and *Delta* subtypes. Regulon activity scores derived using the CollecTRI framework were used to assess differential TF activity across transcriptional subtypes. Barplots show the top positively and negatively enriched TFs (ranked by activity score) for each subtype. Bars are coloured by relative activity score (red = positive, blue = negative). For *Delta*, activity was assessed by comparing *Delta*-OCMs directly against *Gamma*-OCMs to identify regulators specific to this sub-lineage. Enriched TFs recapitulate subtype-specific functional programmes: cell cycle/E2F regulation in *Alpha*, inflammatory and stress-responsive regulators in *Beta*, epithelial identity in *Gamma*, and neuronal/vesicle-related regulators in *Delta*.

**Figure S3.**
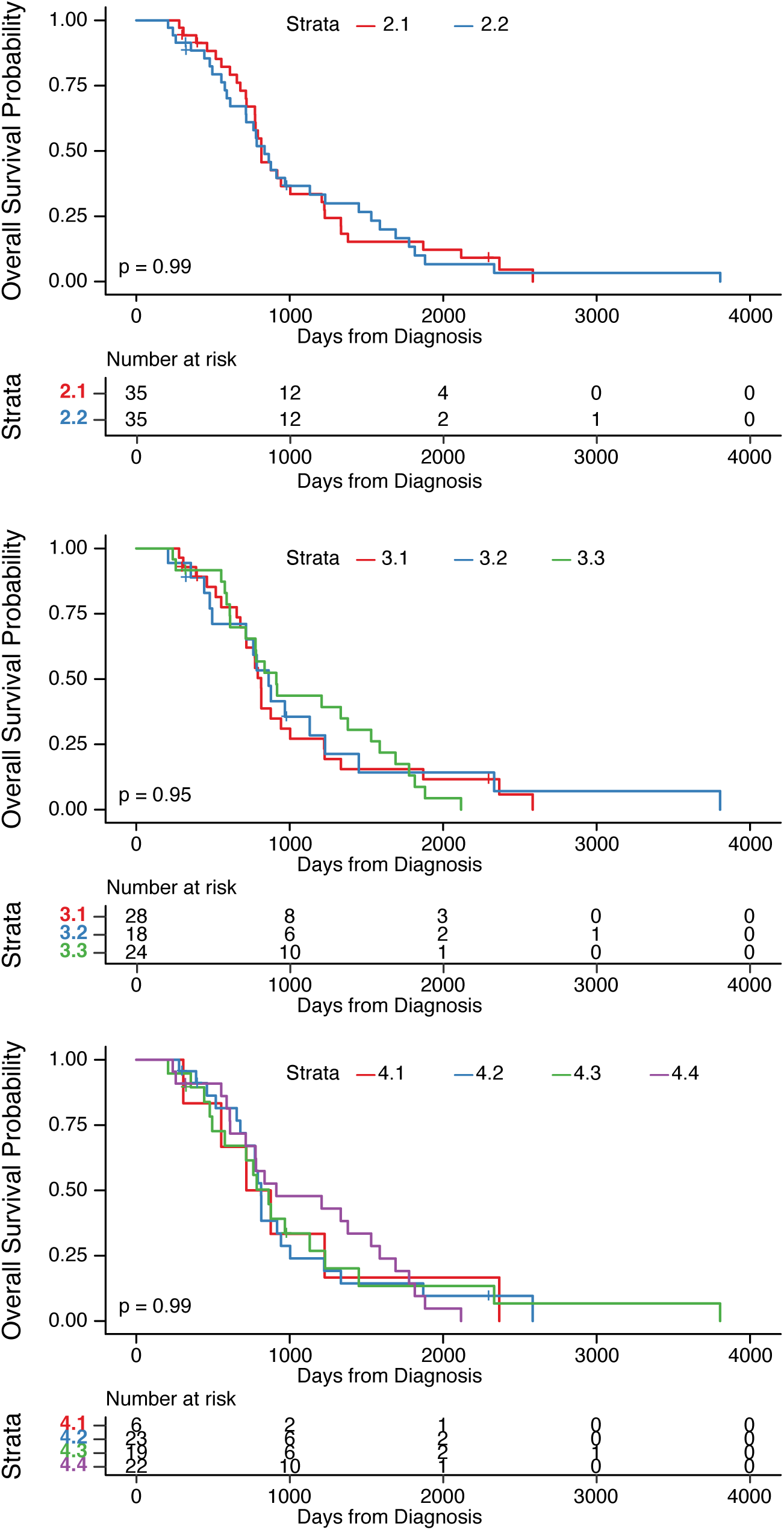
Survival analysis. Kaplan–Meier curves showing overall survival from diagnosis for patients assigned to each transcriptional subtype. Comparisons were performed using the log-rank test. Top panel: Two-cluster solution (2.1 = red, 2.2 = blue). Middle panel: Three-cluster solution (3.1 = red, 3.2 = green, 3.3 = blue). Bottom panel: Four-cluster solution (4.1 = red, 4.2 = blue, 4.3 = green, 4.4 = purple). Tables below each plot show the number of patients at risk at successive time intervals. P-values reflect log-rank test comparisons across the stratified subgroups.

**Figure S4.**
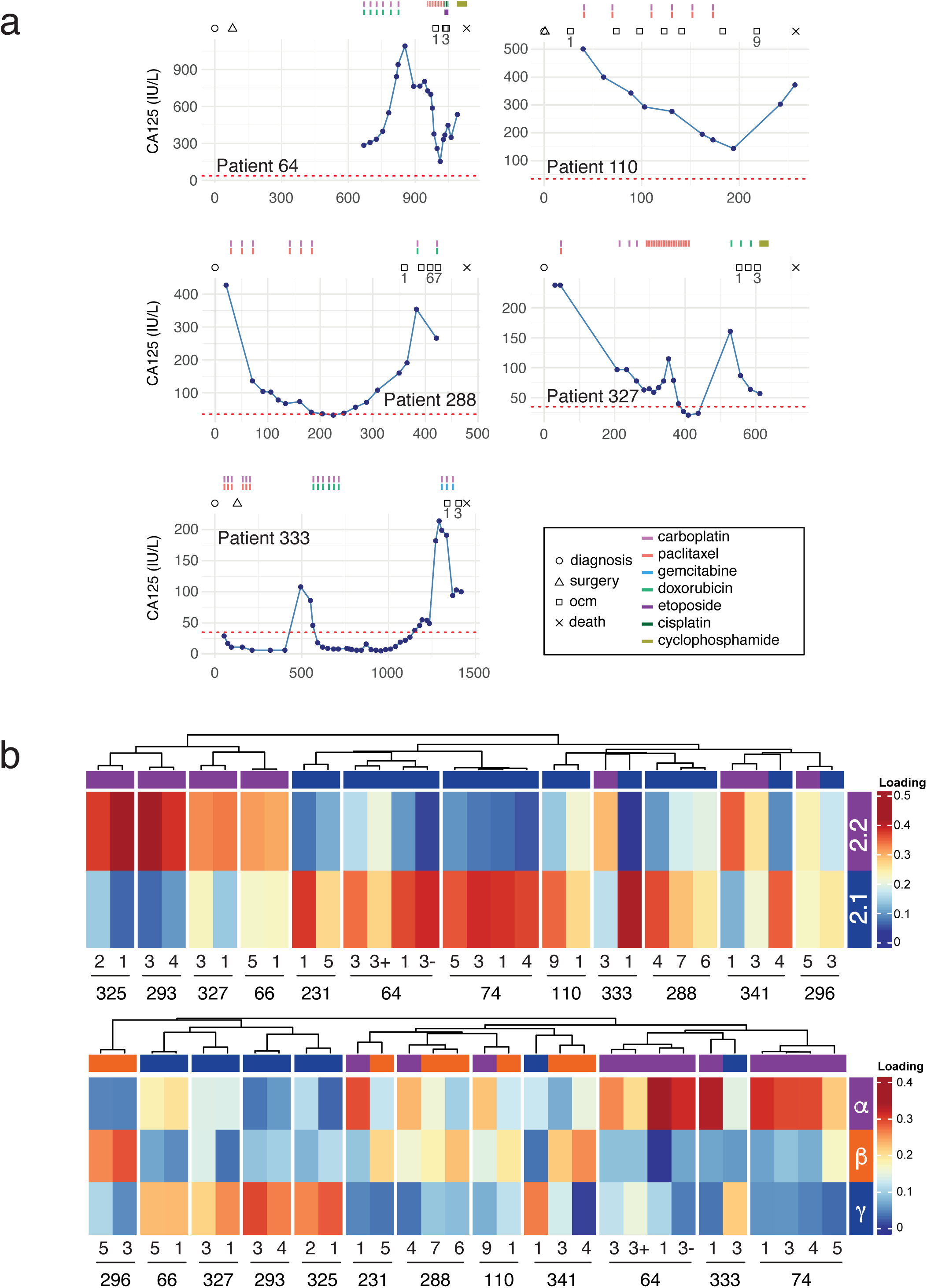
Temporal Resolution of Transcriptional Subtypes. (a) Longitudinal clinical timelines for five additional patients (64, 110, 288, 327, 333), showing CA125 levels over time (blue line) and annotated with diagnosis (circle), OCM sampling (squares), surgery (triangle), and death (cross). Coloured bars below indicate timing of chemotherapeutic exposure: purple = carboplatin, red = paclitaxel, bright green = doxorubicin, blue = gemcitabine, olive green = cyclophosphamide, dark green = cisplatin, and dark purple = etoposide. (b) Non-negative least squares projection of longitudinal OCM samples into the 2-cluster (top) and 3-cluster (bottom) NMF transcriptional spaces. Rows represent clusters, and columns represent OCM samples split by patient number and hierarchically clustered. Heatmap colour indicates loading strength (red = high; yellow/white = intermediate; blue = low). Bar above each column denotes the cluster with the highest loading for that sample, with sample ID and sub-sample numbering shown below.

**Figure S5.**
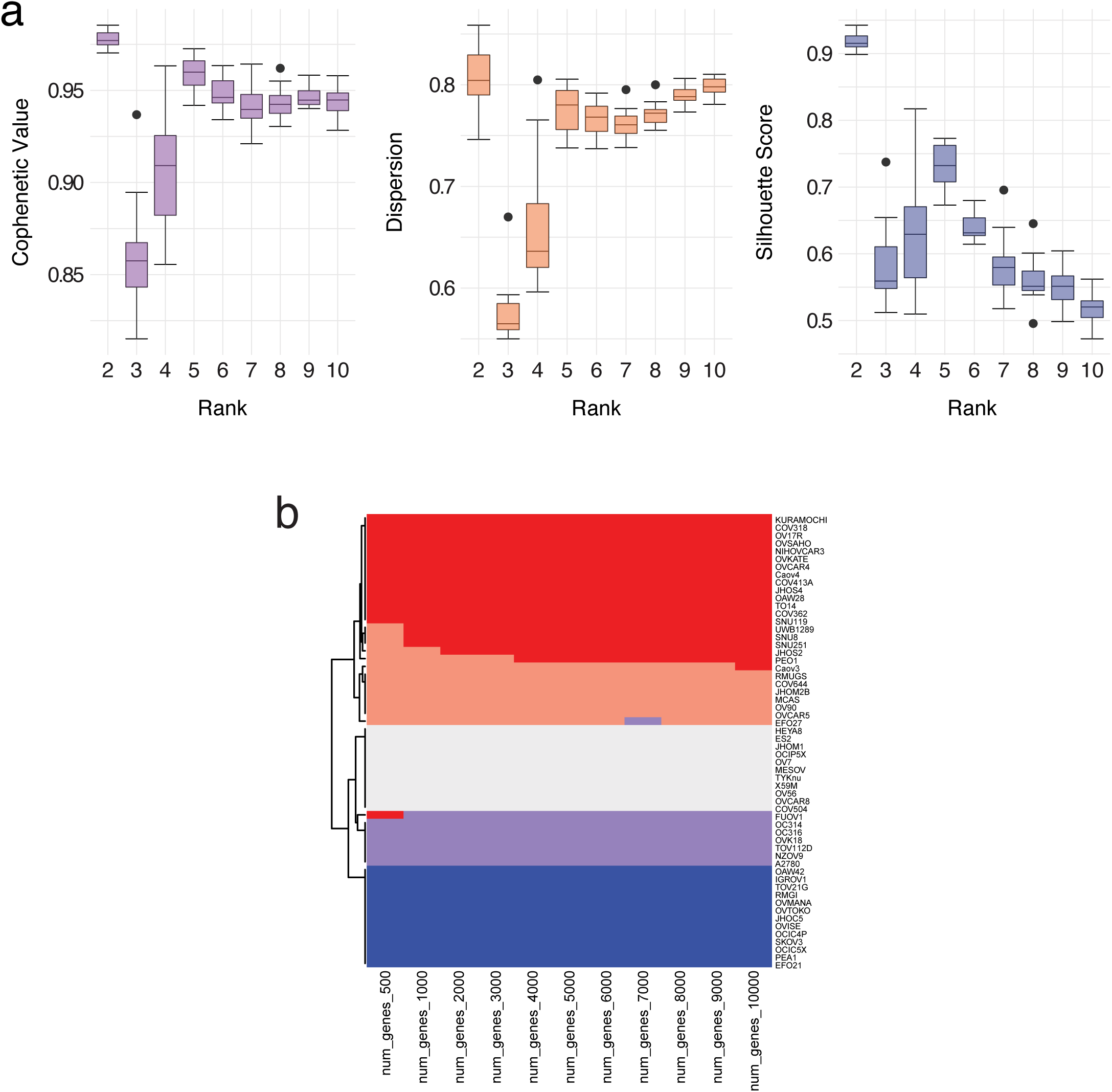
NMF clustering performance and consistency in CCLE cell lines. (a) Evaluation of NMF clustering performance across gene list sizes and cluster ranks. NMF clustering was applied to RNA-seq data from epithelial ovarian cancer cell lines from the Cancer Cell Line Encyclopaedia (CCLE) using gene sets comprising the top 500 to 10,000 most variable genes. For each gene set, clustering was performed across ranks *k* = 2 to *k* = 10 (50 runs per condition). Clustering quality was evaluated using three metrics: cophenetic correlation (left), dispersion coefficient (middle), and silhouette score (right). Each boxplot summarises metric values across the 11 gene sets. (b) Sample-wise cluster assignments across gene list sizes for *k* = 5. Rows represent individual cell lines; columns represent input gene lists (500 to 10,000 genes). Colours indicate assigned cluster identity.

**Figure S6.**
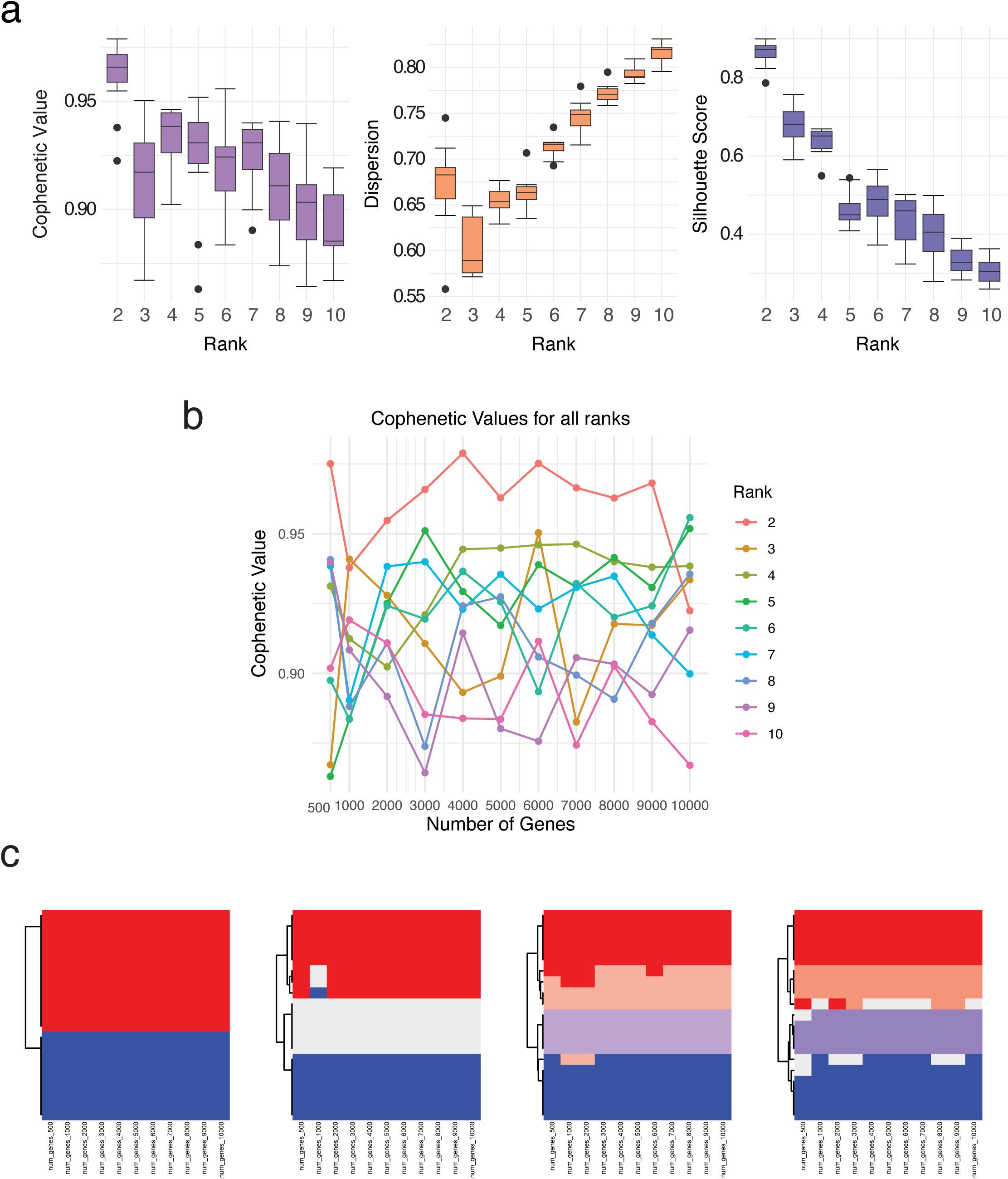
NMF clustering performance and consistency across HGSOC cell lines. (a) Evaluation of NMF clustering performance across gene list sizes and cluster ranks. NMF clustering was applied to RNA-seq data from HGSOC-predicted cell lines from the Cancer Cell Line Encyclopaedia (CCLE) using gene sets comprising the top 100 to 10,000 most variable genes. For each gene set, clustering was performed across ranks *k* = 2 to *k* = 10 (50 runs per condition). Clustering quality was evaluated using three metrics: cophenetic correlation (left), dispersion coefficient (middle), and silhouette score (right). Each boxplot summarises metric values across the ten gene sets. (b) Cophenetic correlation coefficient across input gene list sizes (500 to 10,000 genes) and NMF ranks (, = 2 to 10). Each line represents one rank. Cophenetic values were calculated for 50 NMF iterations per condition and used to evaluate clustering stability. (c) Sample-wise cluster assignments across gene list sizes for *k* = 2 to *k* = 5 (left to right). Rows represent individual HGSOC cell lines (n = 19); columns represent input gene lists (500 to 10,000 genes). Colours indicate assigned cluster identity.

